# Coordinated human prefrontal dynamics sustain task-state representations during learning

**DOI:** 10.64898/2026.05.03.722562

**Authors:** C. Maher, S. Qasim, G. Tostaeva, L. Nunez Martinez, S. Ghatan, F. Panov, A. Radulescu, I. Saez

**Author notes:** Denotes co-senior authorship (A. Radulescu, I. Saez).

## Abstract

Making decisions in complex, real-world environments is challenging. Biologically plausible strategies like reinforcement learning (RL) require attention toward reward-predictive stimuli to define task states, yet how attention and decision processes coordinate in the human brain remains unclear. We hypothesized this arises through interactions between orbitofrontal (OFC) value-based mechanisms and lateral prefrontal (LPFC) attention filtering. To test this, we combined behavioral modeling with local field potential (LFP) and single-unit recordings in 22 subjects performing a multidimensional RL task. Reward expectations were encoded in OFC and LPFC, as reflected in high-frequency LFP and OFC single-unit spiking, but modulated by attention only in LPFC. Theta LFPs encoded reward expectations and indexed attention-dependent LPFC-OFC coordination, with value-related coupling emerging pre-choice in high-attention subjects and post-choice in low-attention subjects. These findings show that prefrontal circuits dynamically coordinate to encode attention-weighted value signals, shaping state representations and providing a tractable solution to learning in complex environments.

## Introduction

Real-world decisions unfold in complex environments containing mixtures of relevant and irrelevant features (e.g., evaluating price, location and reviews when booking a hotel). Exhaustively learning over all possible feature combinations, a strategy often assumed by standard reinforcement learning (RL) frameworks^1,2^, is computationally intractable. A potential solution is to retain only task-relevant information^3–5^, reducing task features into low-dimensional mental models, or state representations. Selective attention acts as a top-down computational process that prioritizes behaviorally relevant features and suppresses irrelevant ones^6^. During learning, selective attention is thought to provide a mechanism for dimensionality reduction by maintaining a compressed, task-relevant state space that supports efficient RL ^3–5, 7–9^. However, how attention interacts with ongoing learning to sustain compressed task states, and how this process is implemented in the human brain, remains unclear.

The prefrontal cortex is well positioned to maintain task-relevant state representations, with growing evidence indicating that flexible cognition arises from distributed network interactions rather than isolated regional computations^10,11^. The lateral prefrontal cortex (LPFC) contributes to top-down attentional control and task-set maintenance^12–15^, enabling dynamic selection and prioritization of behaviorally relevant information. In contrast, the orbitofrontal cortex (OFC) encodes value-based predictions^10,16–19^ alongside abstract task variables that link state representations to expected outcomes^20–23^. Together, these regions support attention and value learning, complementary functions necessary for maintaining efficient task representations.

These observations suggest coordinated interactions between LPFC-mediated attentional control and OFC value representations could support maintenance of compressed task states. We hypothesize that these processes rely on local computation, cross-areal coordination, and integration across OFC and LPFC. More specifically, we postulate that interplay between local high-frequency activity and oscillatory synchronization, which can flexibly couple local computations with inter-regional communication, acts to link PFC attention^7,8,14,24^ with OFC value representations^21,25^.

To test these hypotheses in the human brain, we recorded LFP and single-unit activity from neurosurgical patients performing a multidimensional RL task requiring attention to an instructed feature dimension (shape or color). Behavioral modeling revealed variability in selective attention that predicted performance. In OFC and LPFC, the expected value (EV) of chosen stimuli was encoded in both high-frequency activity (HFA; 60-200 Hz) and theta (3-8 Hz), with distinct temporal profiles. LPFC, but not OFC, exhibited attention-dependent modulation of HFA-EV encoding. LPFC-OFC theta connectivity showed attention-shaped value-dependent coordination: it emerged pre-choice in high-attention participants and post-choice in low-attention participants. Finally, OFC single-units showed rich task coding, indicating neuronal representation of task-relevant information. Together, these findings indicate that attention shapes local prefrontal value signals, while oscillatory synchronization coordinates their interaction to maintain compressed task states during learning in complex environments.

## Results

We administered a multidimensional RL task to 22 neurosurgical patients (age = 39.2 ± 13.8 (M ± SD); sex = 68.2% female; Supplementary Table 1) undergoing monitoring via iEEG for drug-resistant epilepsy (Fig. 1A). Given prior evidence implicating the OFC and LPFC in state representation^21,25,26^, we focused our analyses on these regions. All patients had at least one macroelectrode in both OFC and LPFC (Fig. 1B/C; Supplementary Table 2). During task performance, we recorded LFPs from these regions (Fig. 1B/C). After bipolar re-referencing, we obtained 163 OFC (7.4 ± 4.5; M ± SD) and 122 LPFC (5.5 ± 4.4; M ± SD) bipolar derivations (Fig. 1B/C; Supplementary Table 2). In a subset of five patients, we also recorded OFC single-unit activity using microwires (Supplementary Fig. 1A). Following manual spike sorting and firing-rate thresholding (see Methods; Supplementary Fig. 1B), we identified and analyzed 72 putative single-units (14.4 ± 5.03; M ± SD units/subject).

**Figure 1:**
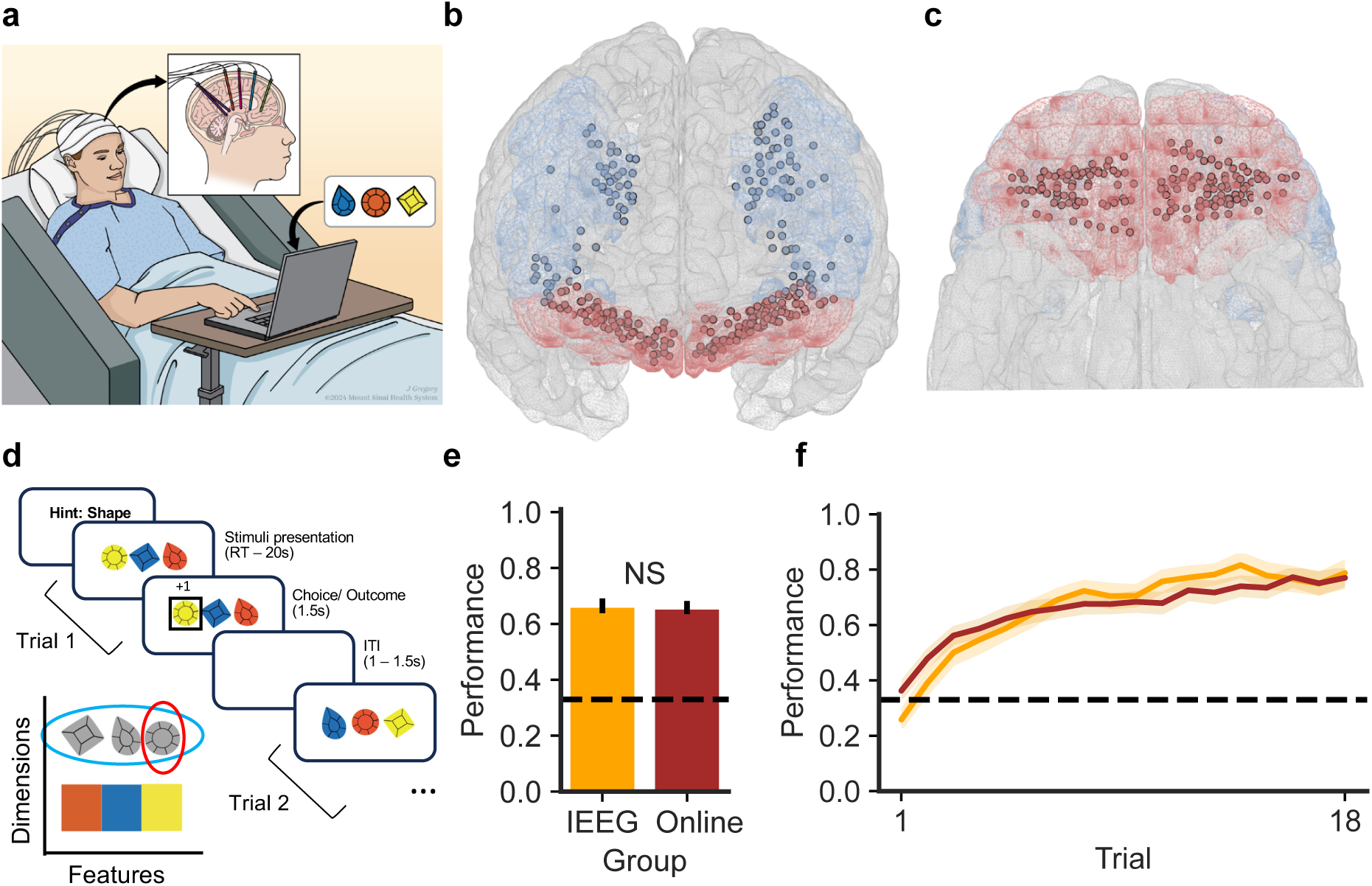
Multidimensional RL task during iEEG recording. **A.** Neurosurgical patients with treatment-resistant epilepsy (N=22) completed the task in the epilepsy monitoring unit while neuronal activity was recorded via iEEG. **B., C.** Macroelectrodes recorded LFPs from the LPFC (blue, N=122 macroelectrodes) and OFC (red, N=163 macroelectrodes). In a subset of patients, we also carried out single-unit recordings. **D.** Multidimensional RL task (Gem Hunters): patients chose between three options that varied in shape and color. On each block (18 trials/block), there was one relevant dimension (e.g., “shape”) and within it one target feature (e.g., “circle”) whose selection maximized the likelihood of reward (80/20 reward probability when the target feature was selected versus a non-target feature). At the start of each block (6 blocks total), patients were told the relevant dimension. Their goal was to learn which feature was the target via trial-and-error. **E.** Behavioral performance (proportion of correct trials) was comparable between iEEG (orange; N=22) and online (maroon; N=51) participants (*t* = 0.10, *p* > 0.05; NS=not significant). **F.** Learning curves show a gradual increase in performance across blocks in both the iEEG and online cohorts, indicating that participants in both groups learned the target feature in a similar manner over time (iEEG: orange; N=22; online: maroon; N=51; shaded area = SEM).

Participants played Gem Hunters, a three-armed probabilistic reward paradigm gamified from prior work^7–9,27,28^ that investigated how attention shapes learning in multidimensional environments (Fig. 1D; see Methods). On each trial, participants chose among three stimuli varying in shape and color. Within each block, they were instructed on the relevant dimension and learned the target feature via trial and error. To ensure performance was not confounded by their epilepsy diagnosis, we validated the task in an independent online sample (N=51; age M ± SD = 35.4 ± 10.18; 31 F). Both groups performed well above chance (online: 65% ± 16%; iEEG: 66% ± 12% correct; Fig. 1E/F) and did not differ (t = 0.10, p > 0.05), confirming successful learning.

### Selective attention modulates behavior during multidimensional RL

Although choices, outcomes, and stimuli are observable, the underlying value learning and attentional processes that we hypothesize are central to task behavior are latent. To infer these computations, we applied behavioral modeling to generate model-derived cognitive variables that could be linked directly to neural activity^29^. To quantify the contribution of top-down selective attention to multidimensional RL, we developed a Selective Attention RL (SA-RL) model that extends feature-based RL frameworks in which agents track the value of individual stimulus features^7–9^ (see Methods). In multidimensional environments, learning becomes tractable when agents prioritize task-relevant features and suppress irrelevant ones, thereby maintaining an efficient state representation^3,4^. Given that even in cases in which participants are explicitly told what to attend to, filtering is imperfect^30^, the SA-RL model formalizes this process by introducing a static, subject-level attention parameter (Φ) that biases both choice valuation and value updating toward the instructed relevant dimension. Fitting Φ as a subject-level parameter captures stable individual differences in selective attention. When Φ approaches 1, agents selectively attend to and learn from the relevant dimension; as Φ decreases toward 0.5, attention and learning are increasingly distributed across both relevant and irrelevant dimensions (Fig. 2A).

**Figure 2:**
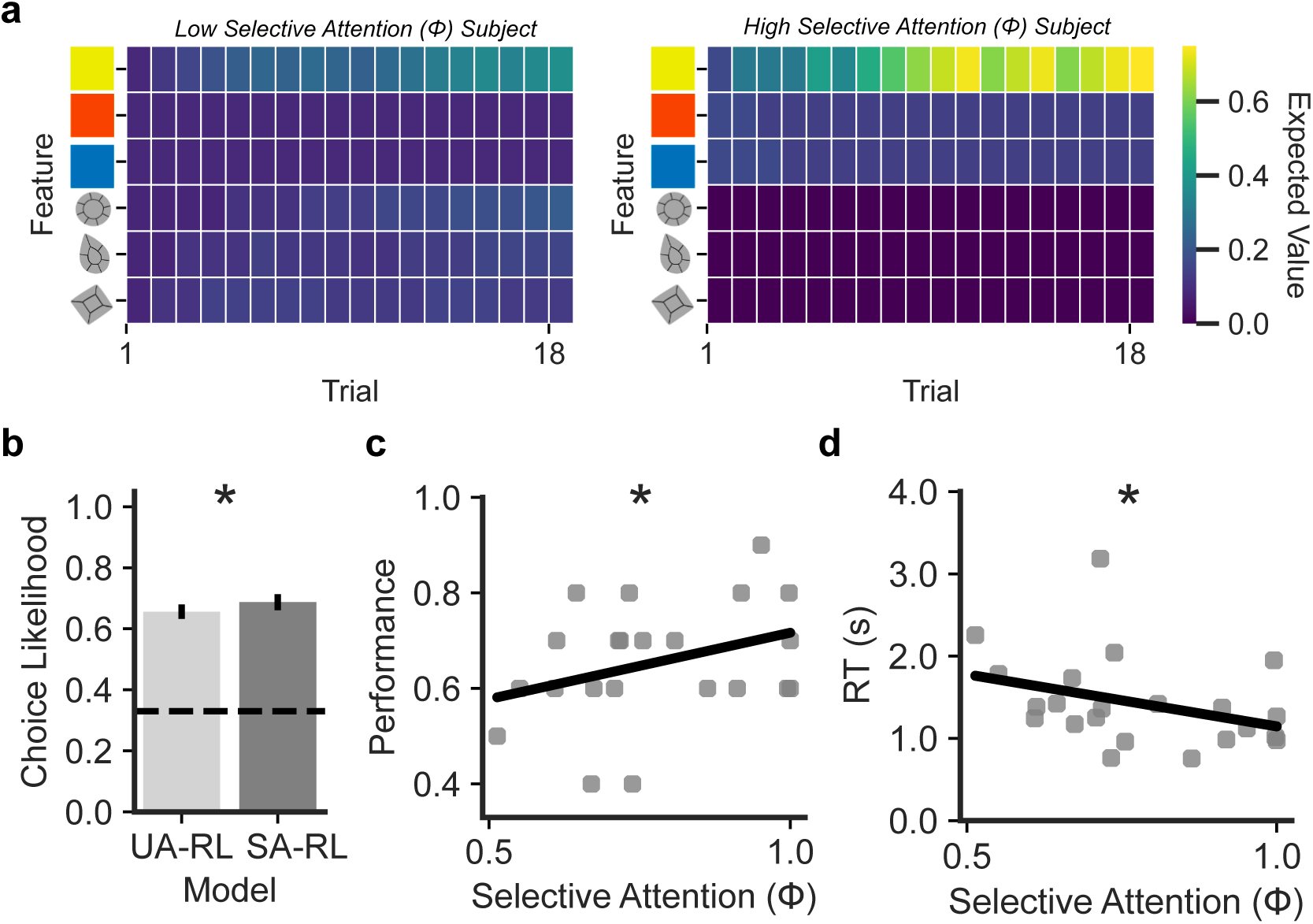
Computational cognitive modeling of choice behavior. Participant choices were fit with two feature-based RL models: a uniform attention model (UA-RL), in which attention was distributed equally across all features, and a selective attention model (SA-RL), in which a subject-level selective attention parameter (ϕ; bounded between 0.51 and 1) biased choice and value updates toward the instructed relevant dimension. Higher values of ϕ indicate stronger selective attention to the instructed relevant dimension. **A.** Example participants illustrating the impact of selective attention on state representation. Each trial depicts the participant’s inferred state representation over environmental features used for value learning. A participant with low attentional selectivity (ϕ = 0.51; left) exhibited diffuse value updates across both relevant and irrelevant feature dimensions. In contrast, a participant with high attentional selectivity (ϕ = 0.99; right) selectively maintains and updates value representations for features within the instructed relevant dimension (e.g., “color”). B. SA-RL explained patient behavior better than UA-RL (t=2.56, p<0.05; error bars = SEM). The SA-RL model’s selective attention parameter was correlated with **C.** task performance (r = 0.39, p < 0.05) and **D.** reaction time (r = –0.44, p < 0.05).

We compared the SA-RL model to a Uniform Attention RL (UA-RL) model, which omits the selective attention parameter (Φ) and weights both dimensions equally during valuation and updating (equivalent to Φ=0.5; see Methods). After parameter and model recovery (Supplementary Fig. 2), we fit both models to each participant’s choices using leave-one-game-out cross-validation and compared fits based on average choice likelihood (see Methods). The SA-RL model explained choices significantly better than the UA-RL model (t= 2.56, *p* < 0.05, paired t-test; Fig. 2B), indicating that patients’ behavior was better predicted using the attention-aware model.

Model fitting yielded subject-specific Φ values (M ± SD = 0.78 ± 0.15; Supplementary Fig. 3), showing individual differences in the degree to which patients deployed selective attention toward the instructed dimension in service of state-space compression. Importantly, these fitted model parameters were associated with model-agnostic task behavior, with higher Φ values associated with better task performance (Spearman r = 0.39, p < 0.05, one-sided; Fig. 2C) and faster reaction times (Spearman r = −0.44, p < 0.05, one-sided; Fig. 2D). These relationships indicate that Φ captures a behaviorally meaningful attentional state, consistent with efficient state representation maintenance (Fig. 2A). The other free parameters in the SA-RL model’s (learning rate and inverse temperature; see Methods) did not correlate with model-agnostic behavioral measures (p > 0.05).

### OFC and LPFC high-frequency and theta activity is modulated during decision-making

We first sought to characterize neural activity during the decision epoch by deriving time-frequency representations (TFRs) of neural activity for each electrode, time-locked to simultaneous choice and reward presentation (decision epoch, t = 0; Fig. 3). Both OFC and LPFC exhibited transient HFA and low frequency modulations, primarily in the theta band (representative single-electrode examples in Fig. 3A/B). These effects were also present at the ROI-level (averages in Fig. 3C/D; significant-shaded averages in Fig. 3E/F; cluster-level p < 0.05). Compared to OFC, LPFC HFA responses were more temporally heterogeneous. Although transient HFA increases were evident at the single-electrode level (Fig. 3B), their timing varied substantially across electrodes, and, unlike in OFC (Fig. 3E), we failed to observe a significant population-level cluster when averaging across LPFC electrodes (Fig. 3F). Band-pass–averaged HFA and theta power for OFC and LPFC, averaged across electrodes within each region (Fig. 3G/H), reveal the temporal profiles of OFC and LPFC activation, highlighting separate volleys of HFA and theta power modulation during pre-choice deliberation and post-choice value updating.

**Figure 3:**
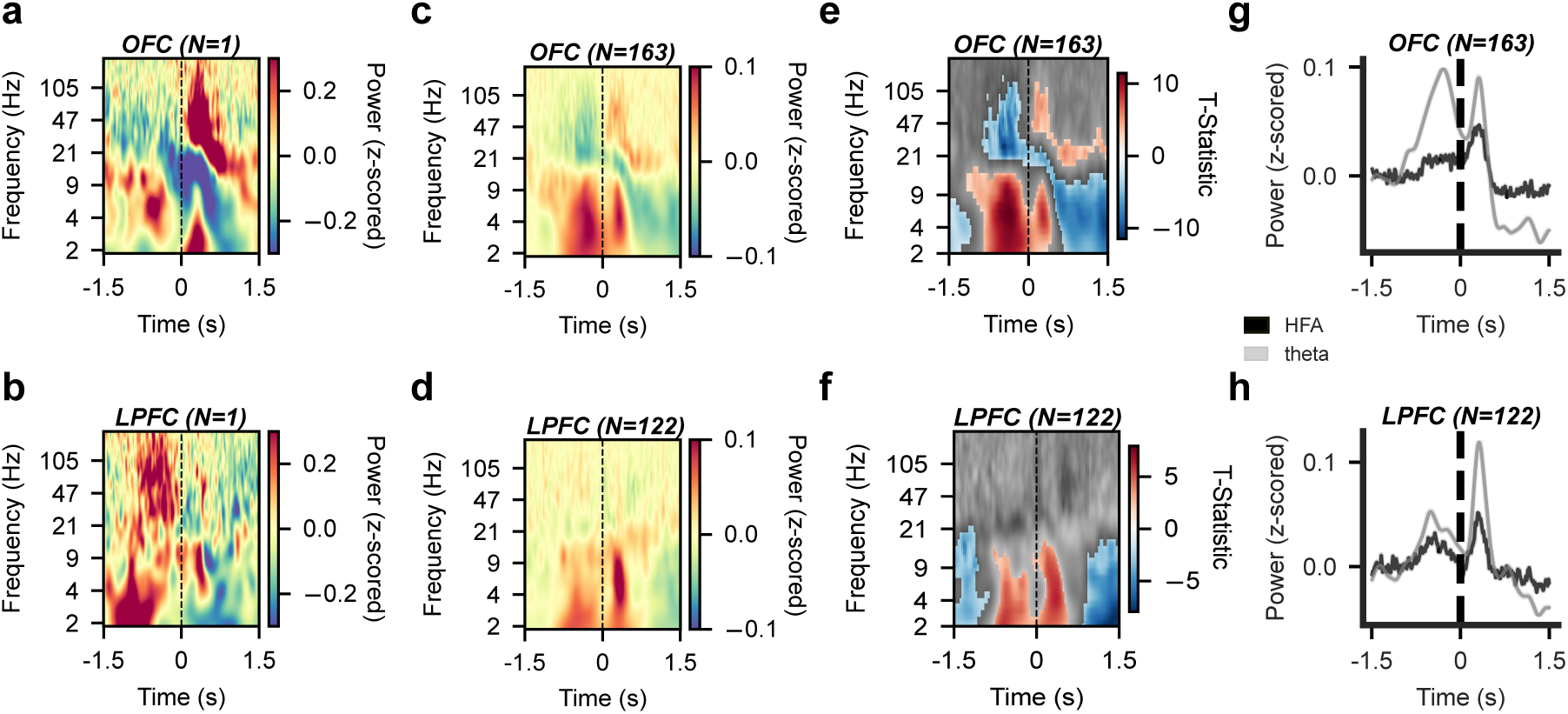
High-frequency and theta activity modulation in OFC and LPFC during decision-making. **A.** Time–frequency representation (TFR) from an example OFC electrode aligned to choice/reward onset (t = 0), illustrating transient, event-locked modulation of high-frequency activity (HFA; 60-200 Hz) and theta (3-8 Hz). **B.** TFR from an example LPFC electrode showing a similar pattern of HFA and theta modulation. **C– D.** Sample-averaged TFRs across all **C.** OFC (N=163) and **D.** LPFC (N=122) electrodes, demonstrating consistent decision epoch modulation of low frequency oscillations in both regions. **E-F.** Colored regions denote clusters of significant power modulation relative to baseline, identified using a one-sample cluster-based permutation test (cluster-level *p* < 0.05). **E.** OFC and **F.** LPFC electrodes. In both regions, significant clusters were observed predominantly at low frequencies, including the theta band (3–8 Hz; cluster level *p* < 0.05). HFA is transient and non-oscillatory; as a result, variability in the timing and expression of these signals across electrodes leads to attenuation when activity is averaged across the sample. **G-H.** Band-averaged power time courses for HFA and theta, averaged across trials and electrodes in **G.** OFC and **H.** LPFC. Time locked fluctuations in both frequency bands indicate coordinated modulation of neural dynamics during the decision epoch in both regions.

Together, these results show the presence of task-evoked activity in both OFC and LPFC around decision, predominantly in HFA and theta, emerging as temporally distinct volleys during pre-choice and post-choice value updating.

### OFC and LPFC activity encode expected value of choice

We next examined how these task-evoked HFA and theta modulations (Fig. 3; Supplementary Fig. 4) related to behavioral computations (Fig. 2). Using the SA-RL model, we derived trial-by-trial regressors corresponding to cognitive variables, namely the expected value (EV) of the chosen stimulus, and used these as regressors to explore the information content of power modulation in pre- and post-choice epochs. EV reflects the learned value of an option over time based on prior outcomes and reflects an internal estimate of reward that is conditional on the maintained state representation, making it a direct readout of how attention shapes valuation. We validated our choice of EV as the main regressor of interest by comparing model fit with two other candidate value signals, namely reward outcome, and reward prediction error (Supplementary Table 3; see Methods). These analyses showed that EV best fit the neural data.

ROI-level TFRs revealed EV-dependent differences in theta and HFA (Fig. 4A), with higher EV associated with increased theta and reduced HFA. To quantify this, we applied a trial-resolved multivariate encoding model relating EV to HFA/theta while controlling for chosen features and relevant dimension (see Methods). At the regional level, both OFC and LPFC showed robust EV encoding in both bands, with higher EV consistently associated with reduced HFA (OFC: β = −1.15, z = −5.19, *p <* 0.001; LPFC: β = −1.42, z = −3.53, *p <* 0.001; Fig. 4B), and increased theta power (OFC: β = 1.53, *z* = 5.71, *p* < 0.001; LPFC: β = 0.96, *z* = 3.21, *p* < 0.001; Fig. 4D), consistent with our previous visual observation (Fig. 4A). At the individual electrode level, we found that a substantial proportion of electrodes encoded EV (HFA: OFC = 40%; LPFC = 49%; theta: OFC = 44%; LPFC = 43%; Supplementary Fig. 4). In contrast, EV-related HFA encoding was absent in a control region, the posterior insula, a region implicated in somatosensory and interoceptive processing^31,32^ but not typically in reward processing (β = -0.35, *z* = -1.05, *p* > 0.05; Supplementary Fig. 6), highlighting a degree of anatomical specificity in EV encoding.

**Figure 4:**
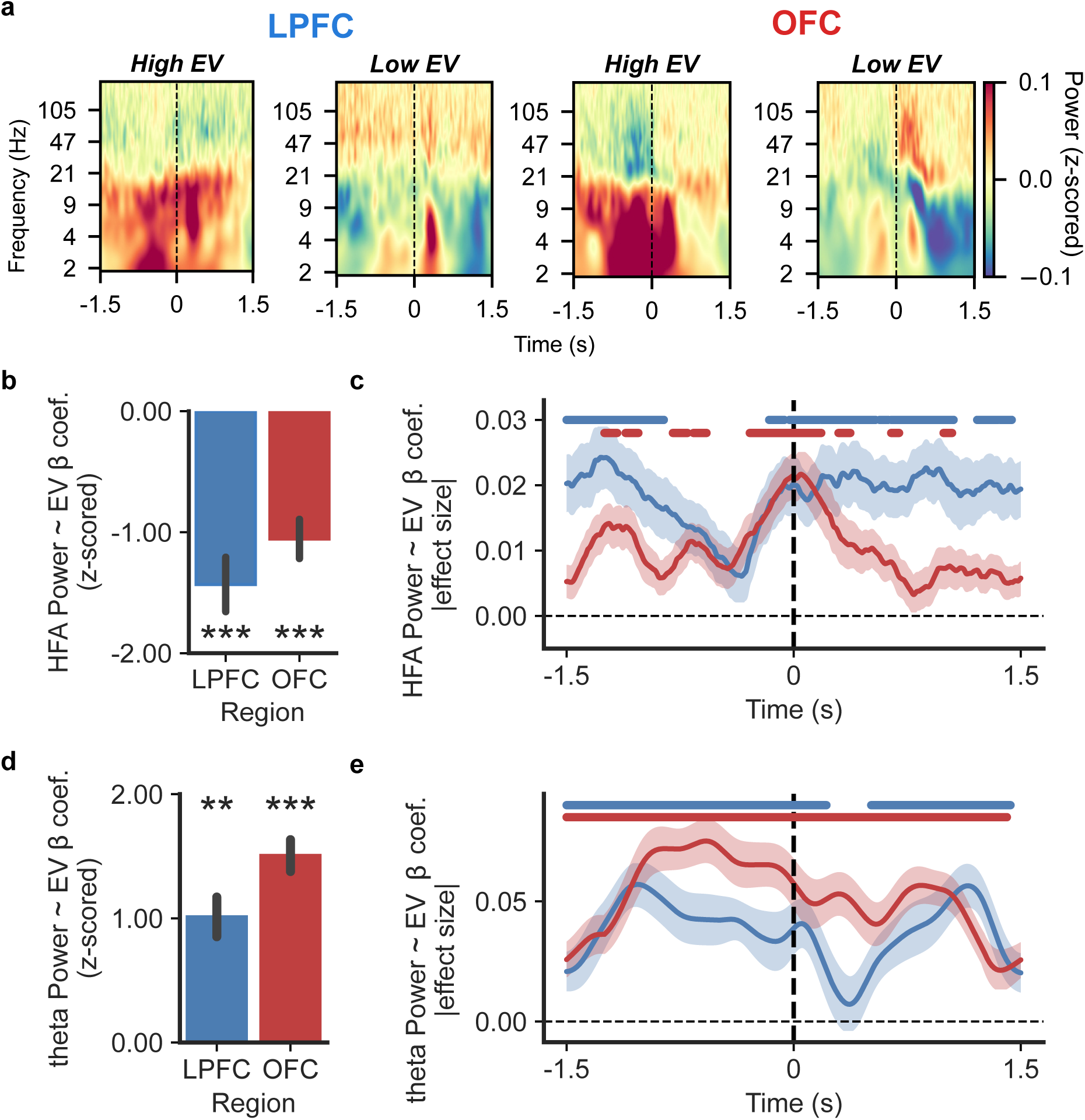
HFA and theta in LPFC and OFC encode EV. **A.** Average time–frequency representations of LPFC (left; N=122 electrodes) and OFC (right; N=163 electrodes) activity reveal differences in activation in both low- and high-frequency bands between high- and low-EV trials. **B.** Both LPFC and OFC exhibit consistent, significant modulation of HFA by EV (*** *p* < 0.001), such that higher EV is associated with reduced HFA across patients in both regions. **C.** Time-resolved HFA EV encoding by region. Absolute EV-related effect sizes from time-resolved regression analyses predicting HFA power, averaged across electrodes within each region (OFC, red; LPFC, blue). Colored lines above each trace indicate time points at which the regional average was significantly greater than zero, as determined by a one-sample cluster-based permutation test (cluster-level *p* < 0.05). OFC HFA EV encoding is more transient, whereas LPFC HFA exhibits a more sustained EV encoding profile across the decision epoch. **D.** Both LPFC and OFC exhibit consistent, significant modulation of theta-band activity by EV (** *p* < 0.01; *** *p* < 0.001), such that higher EV is associated with increased theta power across patients and electrodes. **E.** Time-resolved theta EV encoding by region. Absolute EV-related effect sizes from time-resolved regression analyses predicting theta power, averaged across electrodes within each region (OFC, red; LPFC, blue). Colored lines above each trace indicate time points at which the regional average was significantly greater than zero, as determined by a one-sample cluster-based permutation test (cluster-level *p* < 0.05). Both regions’ theta EV encoding is sustained across the decision epoch.

In contrast to EV-encoding, we did not find group-level evidence for HFA or theta encoding of the perceptual features (relevant dimension and chosen features; all *p* > 0.05), although electrode-level regressions identified a proportion of electrodes that were tuned to relevant dimension (HFA: OFC=19%; LPFC=11%; theta: OFC, 12%; LPFC, 16%), chosen shape (HFA: OFC=19%; LPFC=16%; theta: OFC=18%; LPFC=6%) or chosen color (HFA: OFC=17%; LPFC=13%; theta: OFC=7%; LPFC=19%; Supplementary Fig. 4). Together, these results confirm that observed EV encoding in OFC and LPFC reflects value computations under efficient state maintenance, independent of variability in the state representation’s perceptual content.

We next used time-resolved regression to examine the temporal dynamics of EV encoding (see Methods). HFA profiles differed between OFC and LPFC. OFC showed transient, choice-locked EV encoding, whereas LPFC exhibited sustained pre- and post-choice encoding with a ∼500 ms pause before choice (cluster-level p < 0.05; Fig. 4C). In contrast, theta-band encoding was sustained and similar across regions (cluster-level p < 0.05; Fig. 4E).

Together, these results demonstrate that EV is robustly encoded in both OFC and LPFC in both HFA and theta frequency power, with similar temporal patterns at lower frequencies but different temporal patterns at higher frequencies. This dissociation suggests complementary local encoding dynamics alongside a common low-frequency substrate that may support cross-areal coordination of state representation across the decision process.

### EV encoding in LPFC, but not OFC, is modulated by selective attention

We next asked whether EV encoding was modulated by individual differences in selective attention, as parameterized by the SA-RL model (Supplementary Fig. 3). Specifically, we tested whether individual differences in the selective attention parameter (Φ), which indexes the degree to which the perceptual environment is abstracted into a low-dimensional, task-relevant state representation, predicted the strength of EV encoding (see Methods). We found that selective attention significantly modulated HFA EV encoding in LPFC (*β*= -5.68, *z*= -2.19, *p* < 0.05; Fig. 5A), but not in OFC (*β*= -1.28, *z*= 0.77, *p* > 0.05; Fig. 5E). Specifically, higher selective attention was associated with stronger LPFC HFA modulation by EV, suggesting that local HFA-based EV representation in LPFC was enhanced in individuals who successfully maintained attention during learning. This effect was specific to HFA, and not evident in LPFC or OFC theta (LPFC theta: *β*= -0.07, *z*= -0.23, *p* = 0.82, Fig. 5C; OFC theta: *β*= -0.25, *z*= -0.89, *p* = 0.37, Fig. 5G).

**Figure 5.**
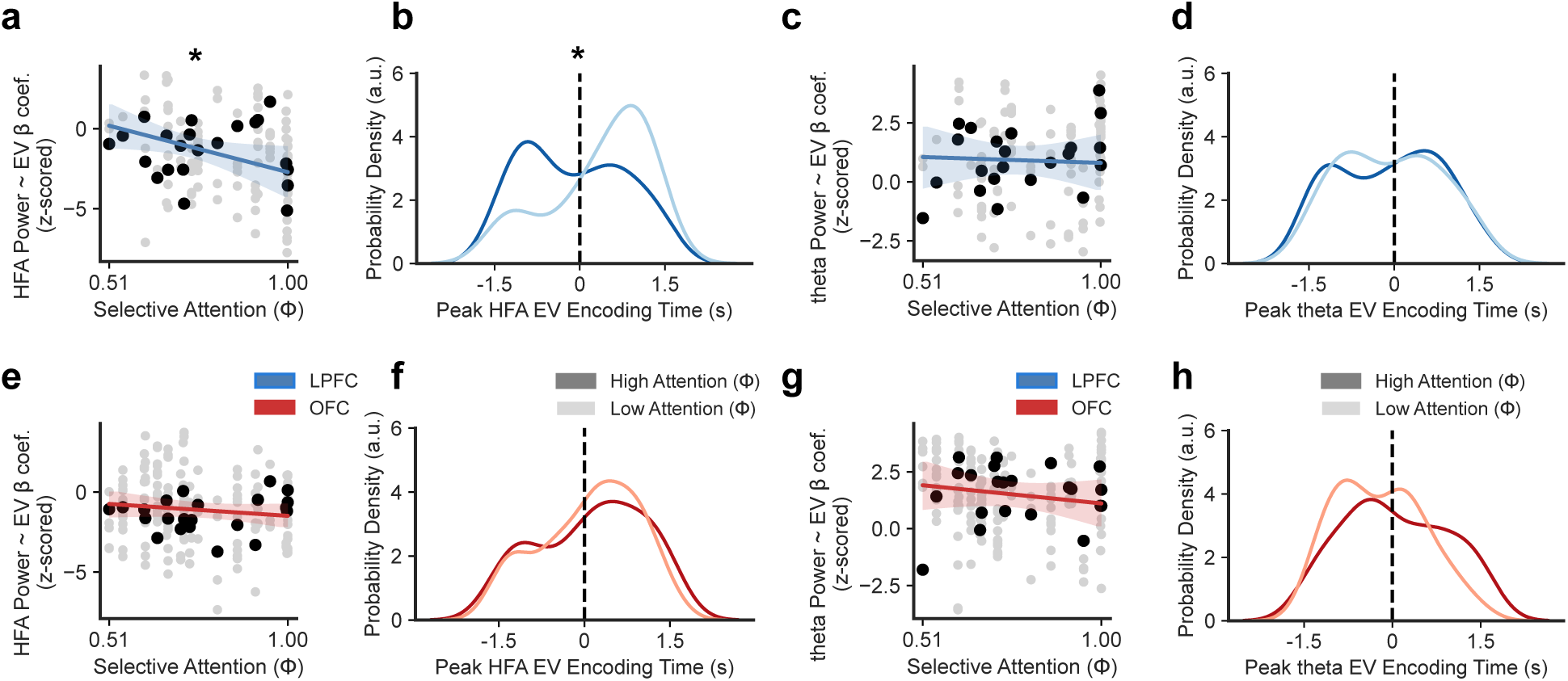
HFA EV encoding is selectively modulated by attention in the LPFC. **A.** Trial-resolved EV encoding magnitude in LPFC HFA was regressed against selective attention weight (Φ) using a mixed-effects model with random intercepts for subject, revealing a significant modulation of HFA value encoding by attention (*β*= -5.68, *z*= -2.19, *p* < 0.05). **B.** To assess temporal effects, we indexed the time point of maximal absolute EV encoding for each LPFC electrode from the time-resolved regression analysis. Electrodes were grouped into high-attention and low-attention groups by a median split of Φ. The resulting distributions showed a significant difference in the peak timing of LPFC HFA-EV encoding between attention groups (KS = 0.28, *p* < 0.05). LPFC encoding peaked prior to choice in high-attention participants (dark blue), and post-choice in low-attention participants (light blue). Median peak latencies were earlier in the high-attention group (median (IQR) = -0.08 s (-0.86 s - 0.62 s)) than in the low-attention group (median (IQR)= 0.63 s (-0.13 s - 1.10 s)). **C–D.** In LPFC, neither the magnitude **C.** nor the peak timing **D.** of EV encoding in the theta band varied significantly as a function of selective attention. **E–H.** Likewise, in OFC, neither the magnitude or timing of EV encoding were varied as a function of selective attention in either **E–F** HFA or **G–H** theta.

LPFC HFA EV encoding’s peak timing differed significantly by attention group, occurring pre-choice in high-attention participants and post-choice in low-attention participants (KS test: D= 0.28, *p* < 0.05; Fig. 5B). In contrast, we found no significant difference in HFA-EV encoding latencies between attention groups in OFC (high-attention group; two-sample KS test: *D* = 0.10, *p* > 0.05; Fig. 5F); or in theta-EV encoding in either region (all *p* > 0.05; Fig. 5D/H).

In summary, we identified attention-dependent modulation of both the magnitude and timing of LPFC HFA, an effect that was region- (LPFC) and frequency- (HFA) specific (Fig. 5). LPFC neural signatures tracked behaviorally derived measures of selective attention, suggesting a role for LPFC in maintaining attention-filtered value representations.

### EV–dependent circuit connectivity is temporally modulated by attention

Cognitive operations involved in state maintenance and value-based decision-making^15,33–35^ are increasingly thought to emerge from coordinated interactions across distributed cortical circuits^33–35^. To test whether OFC and LPFC act in concert to support value-guided state representations, we quantified OFC-LPFC cross-regional functional connectivity using the Phase Slope Index (PSI)^36^, a frequency-resolved measure of directed information flow (see Methods).

Given our initial results (Fig. 3-4) we focused our PSI analyses on theta. For each LPFC–OFC electrode pair, we computed time-resolved theta-band PSI separately for high- and low-EV trials (within-subject mean-based split of z-scored EV values; see Methods). We observed significant EV-dependent modulation of directed connectivity (cluster-level *p* < 0.05; Fig. 6A) in two separate epochs. Prior to choice, LPFC led OFC on high EV trials. Following choice, the pattern reversed, with LPFC leading OFC on low EV trials.

**Figure 6:**
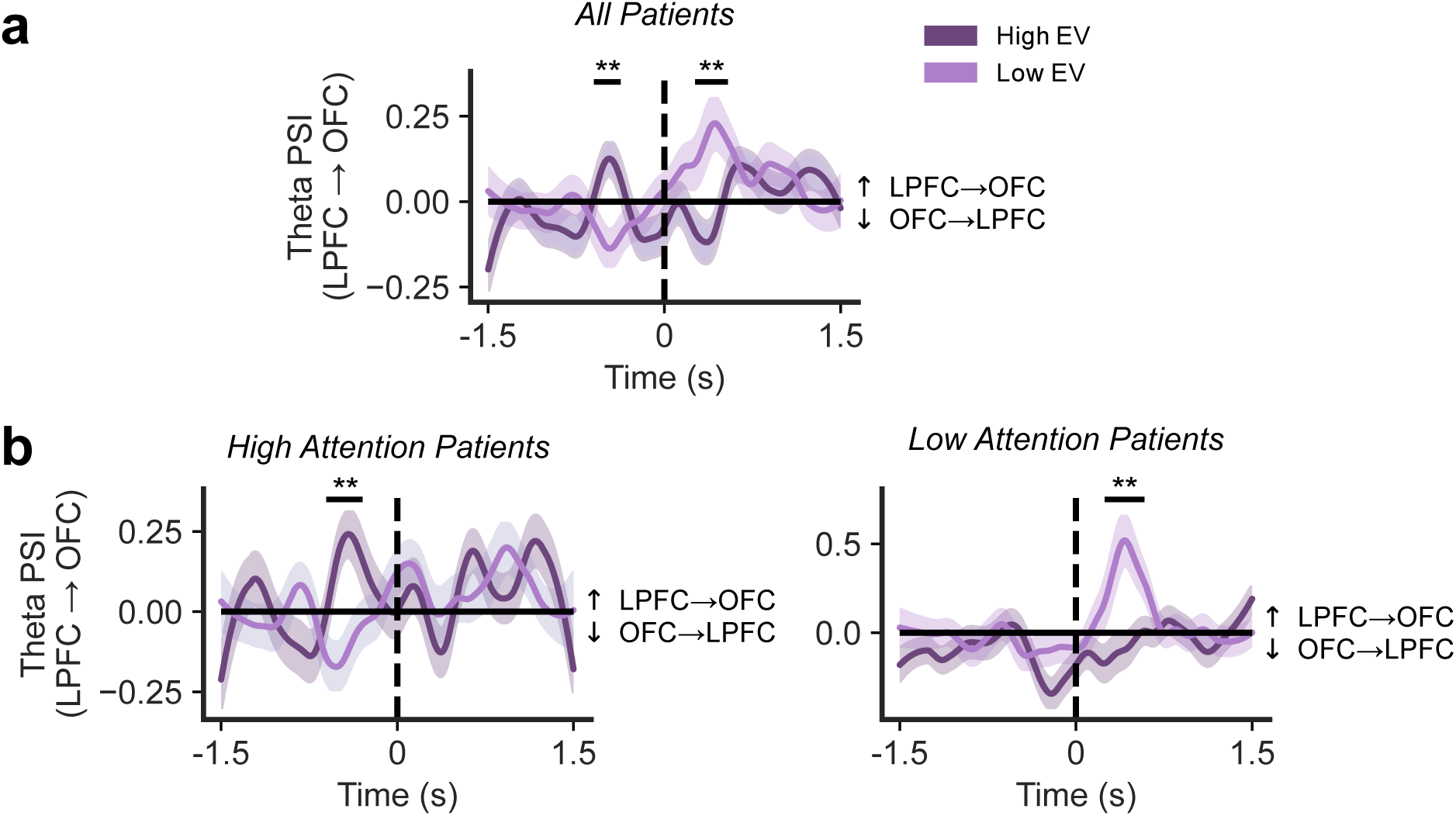
LPFC–OFC theta-band connectivity reflects attention- and value-dependent dynamics. **A.** We examined the time course of Phase Slope Index (PSI), a proxy for directed communication, to examine the nature and timing of circuit-wide interactions between LPFC and OFC. We computed time-resolved theta PSI across all patients with ipsilateral LPFC–OFC electrode pairs (N=22 patients) and compared PSI in high vs. low EV trials at each timepoint across the decision epoch (high EV: within-subject z-scored EV ≥ 0; low EV: < 0). Positive PSI values indicate LPFC leading OFC, while negative values reflect OFC leading LPFC. Two-sample cluster-based permutation test identified two significant epochs (gray shading; cluster level p < 0.05): before and after choice/reward (t=0), in which connectivity differed significantly by value context. Pre-choice LPFC-to-OFC connectivity was stronger on high EV trials, whereas post-choice connectivity was stronger on low EV trials. **B.** Splitting patients into high- (left) and low-(right) attention groups revealed a temporal dissociation in pre- and post-choice functional connectivity epochs by attentional strategy. In high-attention patients (left), the pre-choice LPFC-to-OFC connectivity epoch was preserved, consistent with LPFC attentional control during pre-choice. In low-attention patients (right), only the post-choice OFC-to-LPFC connectivity epoch survived, suggesting the engagement of OFC reward learning mechanisms.

Next, we tested whether these connectivity dynamics between LPFC and OFC were attention-dependent, by repeating this analysis separately in high- and low-attention patient groups (Φ median split, n=11 patients in each group). This analysis revealed attention-dependent differences: high-attention participants showed significant EV-dependent LPFC-OFC connectivity during the pre-choice epoch (cluster-level *p* < 0.05; Fig. 6B), whereas low-attention participants showed significant EV-dependent LPFC-OFC connectivity only post-choice (cluster-level *p* < 0.05; Fig. 6B). Thus, attentional state was associated with a shift in the timing of EV-related LPFC–OFC connectivity, with high selective attention linked to pre-choice LPFC–OFC interactions and lower selective attention linked to post-choice LPFC–OFC interactions.

Together, these results show that theta connectivity, consistent with its role in functional interregional communication^37–40^, coordinates LPFC and OFC activity, with its temporal dynamics shaped by attentional state.

### Local LPFC value signals interact with attentional strategy to predict LPFC-OFC connectivity

We next sought to examine whether a relationship existed between local and circuit levels of state maintenance mechanisms. Specifically, we examined whether the pre-choice attenuation in LPFC HFA–EV encoding (Fig. 4C), which coincides in time with the increase in LPFC–OFC theta connectivity (Fig. 6A; overlaid in Fig. 7A), were related. If so, moment-to-moment variation in LPFC EV encoding should covary with its theta-band coupling to OFC. To test this, we modeled theta connectivity using a mixed-effects model including an interaction between LPFC HFA EV encoding strength (HFA β) and selective attention (Φ) during the pre-choice period, with electrode as a random intercept (see Methods).

**Figure 7:**
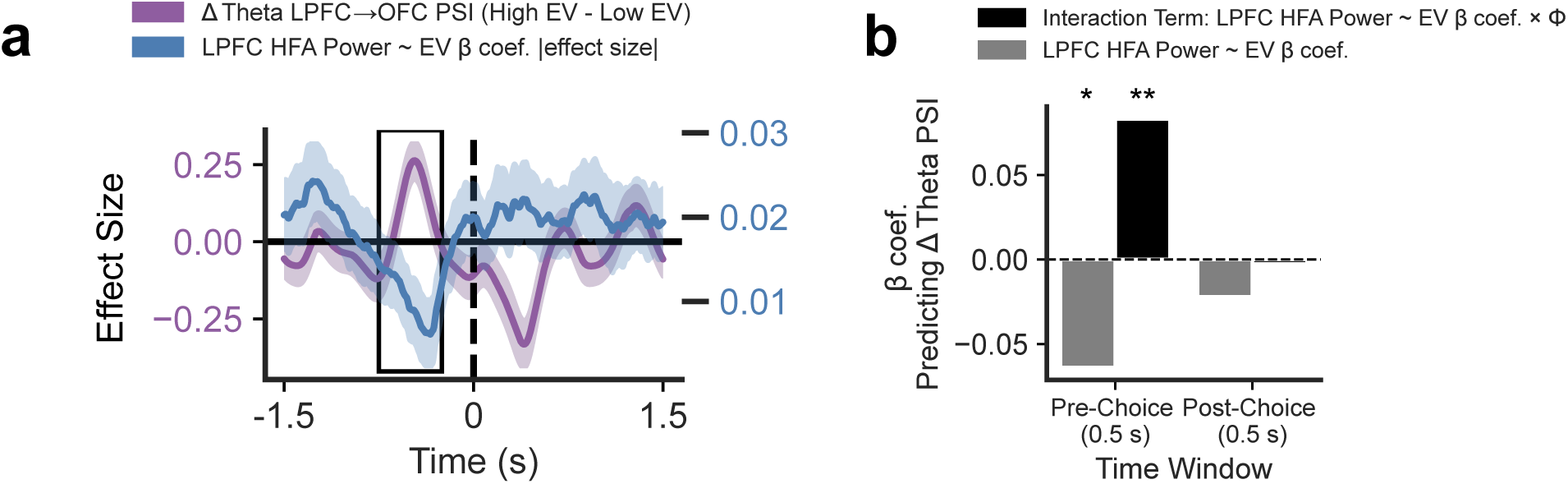
LPFC local value signals modulate LPFC–OFC connectivity in an attention-dependent and time-sensitive manner. **A.** Time courses of the *β* coefficient from the time-resolved HFA-EV regression in LPFC electrodes (blue trace; from Fig. 4C) and the ΔPSI (high–low EV; purple trace; from Fig. 6A). The black square denotes a 500 ms window, which contains the pre-choice significant ΔPSI cluster, during which temporal alignment between local HFA value signals in the LPFC and LPFC-to-OFC PSI occurs. **B.** A multivariate mixed-effects model predicting ΔPSI (high–low EV) from LPFC EV encoding (HFA∼EV β) and its interaction with selective attention weight (Φ) within the pre-choice window (black square; panel A) revealed that LPFC HFA EV encoding significantly predicted LPFC–OFC connectivity dynamics (β = −0.06, z = −2.59, p < 0.05), and this relationship was significantly moderated by selective attention (β = 0.08, z = 2.79, p < 0.01). In a matched post-choice window, LPFC EV encoding showed no significant relationship with ΔPSI, irrespective of attentional strategy, indicating that this relationship is both attention- and time-specific.

During the pre-choice period, we found a significant association between the strength of LPFC EV-encoding and interregional connectivity dynamics (β = −0.06, z = −2.59, *p* < 0.05; Fig. 7B). Notably, the interaction between HFA β and Φ was also significant (β = 0.08, z = 2.79, *p* < 0.01; Fig. 7B), indicating that selective attention not only modulated the magnitude but also the direction of the relationship between local LPFC EV encoding and LPFC-OFC theta-band connectivity. In contrast, during the post-choice epoch neither local LPFC EV encoding (*β* = -0.02, z= -0.69, *p* > 0.05) nor its interaction with attention predicted LPFC-OFC theta connectivity (*β* = -0.003, z= -0.07, *p* > 0.05; Fig. 7B).

Together, these findings suggest a time-specific shift from local value encoding in LPFC to coordinated LPFC–OFC interactions, modulated by individual differences in task-state maintenance (Φ).

### Single-units in OFC encode state-relevant variables with profiles consistent with population-level signals

Finally, we complemented population-level LFP analyses with single-unit recordings to examine task-state representations at the cellular level and across scales. In a subset of patients (N = 5), we recorded from OFC (n = 72 units; Methods; Supplementary Fig. 1A/B). We assessed encoding of EV and the instructed relevant dimension, which together define the compressed task-state.

Applying the same non-parametric LFP encoding model to individual neuron firing rates (see Methods), we found that a substantial proportion of OFC neurons were tuned to EV (28/72, 36%) and instructed relevant dimension (23/72, 39%; both *p* < 0.001 binomial test vs 5% null; Fig. 8B/D). These populations overlapped, with a significantly greater proportion of units encoding both variables vs random mixing (13 units; 34% of task-selective units; binomial test, *p* < 0.001; Supplementary Fig. 1C).

**Figure 8:**
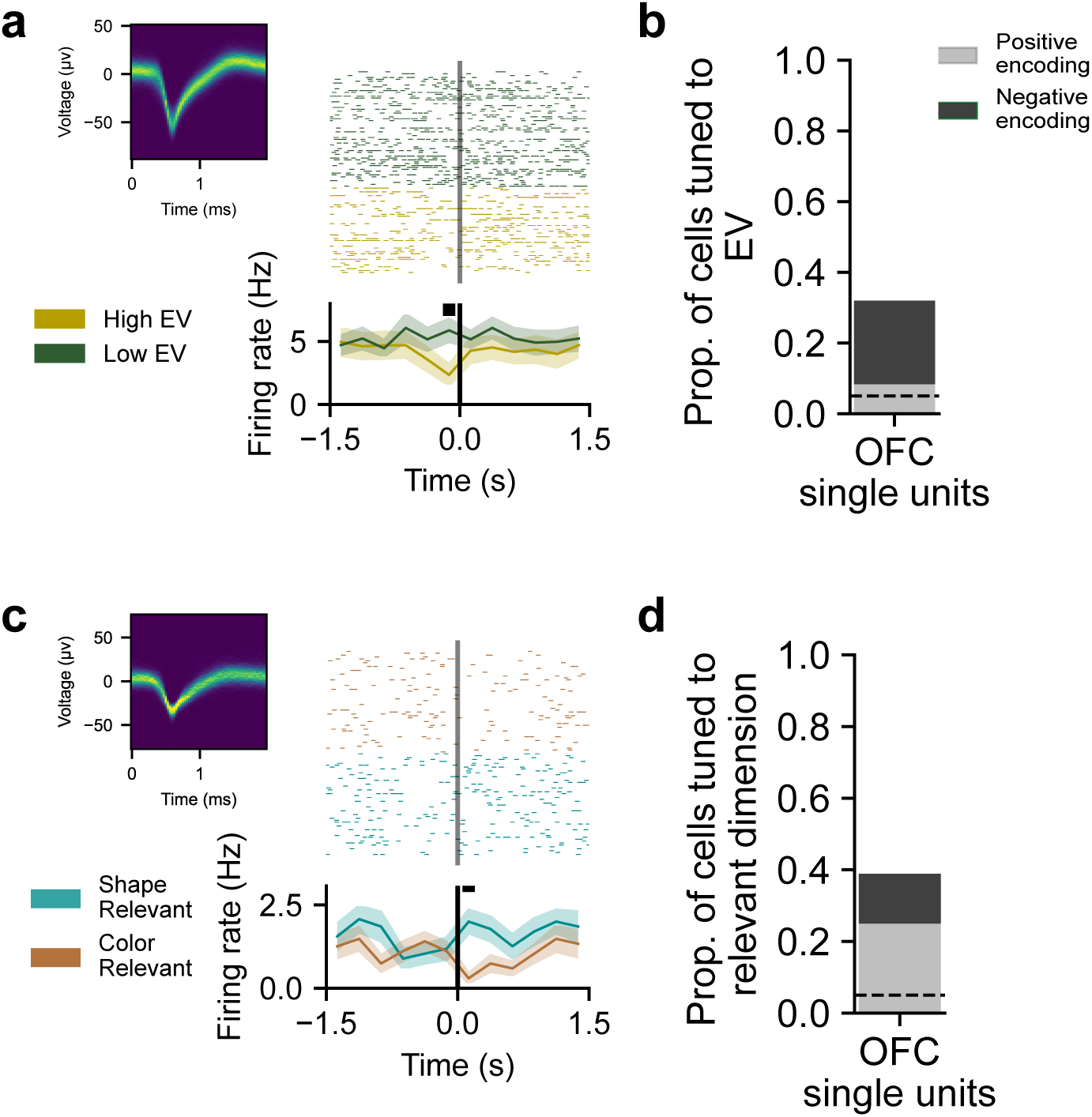
OFC single-units encode state-relevant information. **A.** Raster and peri-event firing rate from an example putative OFC unit showing a significant, time-locked difference in firing rate between high- and low-EV trials during the pre-choice period (*FDR-corrected p* < 0.05, shading = SEM). **B.** Across putative OFC units, a substantial proportion exhibited significant tuning to EV (36%), exceeding chance levels. EV tuning was assessed using a trial-resolved multivariate regression predicting mean firing rate from EV, relevant dimension, and chosen features, with statistical significance determined relative to a permutation-derived null distribution generated by shuffling EV labels. Among units selective for EV, 8% exhibited increased firing with higher EV, whereas 24% exhibited decreased firing, indicating a predominant negative tuning pattern. **C.** Raster and peri-event firing rate from an example putative OFC unit showing significant modulation by the relevant dimension (shape vs. color; *FDR-corrected p* < 0.05). **D.** Across putative units, a well-above-chance proportion exhibited significant tuning to the relevant dimension (38%), assessed using the same regression framework with permutation testing based on shuffled relevant-dimension labels. Among units selective for the relevant dimension, 25% increased their firing rate when shape was relevant (relative to color), whereas 14% showed the opposite pattern, decreasing their firing rate when shape was relevant.

Interestingly, the directionality of single-unit EV and relevant dimension effects (Fig. 8B) mirrored those observed in OFC HFA (Fig. 4B). Increasing EV was associated with a suppression of HFA in both OFC (Fig. 4B). Similarly, most single-units were negatively tuned to EV (24% EV-selective units were negatively tuned; binomial test p<0.05 vs 50/50 split; Fig. 8B).

Single-unit effects for relevant-dimension coding similarly paralleled those observed in OFC HFA. At the population level, a substantial proportion of OFC electrodes exhibited significant HFA modulation by the relevant dimension (19%; Supplementary Fig. 4); however, unlike EV encoding in OFC (see Results: HFA EV encoding), this effect did not exhibit consistent group-level directionality. This pattern indicates heterogeneous encoding of the relevant dimension at the macroelectrode level.

A comparable pattern emerged at the single-unit level. Among units selective for the relevant dimension, 25% increased their firing rate when shape (relative to color) was relevant, whereas 14% decreased their firing rate. A binomial test revealed no significant deviation from a 50/50 distribution of positive versus negative tuning (p > 0.05; Fig. 8D), further supporting the absence of a systematic directional bias.

Together, these findings show that compressed task-state representations are instantiated at the level of single neurons in OFC and expressed at the population level through HFA, bridging cellular and circuit-level account of value-based learning in the prefrontal cortex^41^.

## Discussion

Learning in complex environments requires restricting value learning to the subset of features that matter for the current task^3,4,22,42^. Here, we tested the idea that selective attention solves this problem through coordination of local activity and cross-regional communication between the LPFC and OFC. By combining an attention-aware RL model^7–9,27,28^ with human intracranial recordings, we found that individual differences in attentional filtering predicted behavior and were reflected in distinct local and circuit-level dynamics across LPFC and OFC. These findings refine state-representation accounts of OFC^21,22^ by showing how attention-weighted value signals are dynamically coordinated across prefrontal cortex areas to meet task demands, with LPFC providing top-down attentional filtering.

Our behavioral model captured individual differences in selective attention with a single parameter, Φ, that biased valuation and learning toward the relevant dimension. Higher Φ predicted better performance and faster responses, consistent with attention acting as a computational filter over the RL state space^3,4,8^. Notably, this variability persisted despite explicit task instructions, indicating that attentional strategy during state maintenance differs across individuals, consistent with prior work demonstrating inter-individual variability in the attentional strategies deployed during multidimensional RL^9^.

At the neural level, we found that LPFC and OFC encoded model-derived value estimates in HFA and theta (Fig. 4), with these signals playing distinct and complementary roles. In both regions, higher EV was associated with lower HFA and higher theta (Fig. 4B/D). The inverse relationship between EV and HFA is consistent with cross-species evidence of heterogeneous prefrontal value coding^16,17^ and raises the possibility that, as value becomes more certain, local activity becomes more selective or efficient, leading to reduced broadband power. The observed trade-off between HFA and theta power as a function of value (Fig. 4B/D) suggests that HFA may index local population computations, while theta provides a shared substrate for communication across regions^43–45^. This finding is also consistent with a push-pull relationship between low- and high-frequency activity thought to reflect coordinated shifts between large-scale integrative processes and local spiking related computations^46^.

This interpretation was strengthened by convergent OFC LFP and single-unit activity. Most EV-selective OFC neurons were negatively tuned, with firing rates decreasing as a function of increasing EV (Fig. 8B), mirroring the negative relationship between EV and OFC HFA (Fig. 4B). By contrast, coding of the relevant dimension, which was also present at the cellular level (Fig. 8D), was more heterogeneous, with no difference in the proportion of neurons showing positive versus negative tuning (Fig. 8D). This was in turn reflected by a lack of group-level directionality in HFA (Supplementary Fig. 4). Thus, EV showed relatively homogeneous directionality across scales, whereas state-content information was represented in a more distributed and heterogeneous format which could be obscured at the HFA population level. This pattern is consistent with the idea that global, salient variables (e.g., value) are expressed through broad population-level modulations, whereas task-relevant state information is carried by more heterogeneous neural ensembles^47^. More broadly, the convergence between single-unit firing and HFA supports the interpretation that OFC HFA provides a useful population-level readout of local spiking and postsynaptic activation dynamics^48^ when cellular encoding is homogeneous, but less so when there is diversity in cellular encoding schemes.

Despite shared EV coding (Fig. 4), LPFC and OFC differed in both the temporal structure of these signals and in how strongly they depended on attentional strategy (Fig. 5). OFC HFA encoded EV transiently around decision events, consistent with a role in expressing state-dependent value estimates around the time of choice^16,21,25^, and this encoding was insensitive to attentional biases. LPFC HFA, in contrast, was more sustained across the decision epoch and varied systematically with attention (Φ), such that participants with stronger selective attention showed earlier and stronger LPFC EV signals (Fig. 5A/B). These findings support a division of labor in which LPFC helps constrain which features define the current task state, while OFC represents the value of options, and potentially other task-relevant information, within that attention-weighted state space^5,22^. LPFC EV dynamics appear before and after choice, potentially indexing how the state space is constrained prior to choice evaluation and post-reward outcome^25^ for selection and learning, respectively. LPFC dynamics, in addition, are modulated by attentional state: when attention is focused, value signals are stronger and emerge earlier; when attention is diffuse, they are delayed and less robust.

Beyond this local encoding, our circuit-level analyses show that LPFC and OFC are dynamically coordinated as a function of both value and attentional strategy through theta-band phase relations. Specifically, directed theta connectivity analyses revealed a value-dependent reconfiguration of LPFC-OFC interactions across decision processes. Before choice, high-value trials were associated with stronger LPFC-to-OFC theta coupling, while low-value trials showed relatively stronger OFC-to-LPFC interactions (Fig. 6A), a pattern that was reversed post-choice (Fig. 6A). Consistent with the notion that these directional interactions reflect attentional influences, the pre-choice LPFC-to-OFC value-dependent coupling was driven by high-attention participants, whereas post-choice coupling was primarily expressed in low-attention participants. Together, these observations suggest that prefrontal connectivity is dynamic^33,35,38^, rapidly shifting depending on attentional state and value estimates. When value estimates are stable, LPFC may exert top-down influence over OFC consistent with proactive maintenance of task-relevant information^14,24^, whereas when value is lower or more uncertain, OFC may engage LPFC to support value updating or attentional reallocation^21,25^. Rather than implying a fixed hierarchy, these results favor a dynamic account in which the balance of influence between prefrontal regions changes across decision phases^24,33,49^.

Finally, the observed relationship between local LPFC HFA value signals and LPFC-OFC theta connectivity during the pre-choice period suggests an interplay between local EV encoding and theta circuit dynamics^44^ (Fig. 7). Selective attention altered not only the magnitude of LPFC value coding, but also its relationship to interregional communication, directly linking local computations to distributed network dynamics^43–45^. Together, these findings support the view that selective attention facilitates efficient state maintenance by aligning LPFC value representations with LPFC-OFC coordination at the time those representations are most useful for guiding choice (i.e. during deliberation).

Our study presents several limitations. First, the task isolates maintenance of an instructed relevant dimension, so our results speak most directly to sustaining a compressed task state once the relevant dimensions are known, rather than to discovering latent task structure or switching among candidate state spaces in more naturalistic environments. Second, we modeled attention as a stable subject-level parameter (Φ) to capture robust individual differences, but attentional filtering almost certainly fluctuates across trials and learning stages. A dynamic model of trial-by-trial attention may therefore capture additional variance and help explain when proactive versus reactive control modes emerge. Third, the study was necessarily constrained by clinical sampling. As participants were patients with epilepsy, anatomical coverage was determined by clinical need, and single-unit data were available only in OFC and only in a subset of participants. Finally, directed PSI theta coupling is consistent with coordinated information flow^37,39,40,50^ but does not by itself establish causality, and other mechanisms of cross-regional coordination could be at play. Future work combining dynamic behavioral models with causal perturbations^51^ will be important for refining this circuit account and the causal role of OFC and LPFC.

Together, our findings support a multiscale prefrontal account of how selective attention enables learning in multidimensional environments. Under this account, LPFC helps define the task-relevant state and OFC expresses value within that state^5,22^, with theta-band coordination linking these computations across regions and decision phases^39,40^. More broadly, this work connects computational theories of attention-weighted RL^7,8,27^ to their physiological implementation in the human brain and highlights flexible coordination across prefrontal circuits as a core principle of adaptive behavior^33^. These results provide a cross-scale framework for future work on how state representations are stabilized, updated, and disrupted in conditions marked by impaired cognitive control and maladaptive learning^52^.

## Methods

### Task

Twenty-two patients with drug-resistant epilepsy completed a multidimensional RL task, “Gem Hunters”, during intracranial monitoring. The task was adapted from prior work^7–9,27,28^ and gamified for use with neurosurgical patients. Experimental sessions took place at the patients’ bedsides using a laptop computer. The task was created using JavaScript and custom functions adapted from jsPsych^53^. All procedures were approved by the Institutional Review Board of the Icahn School of Medicine at Mount Sinai (New York, NY), and all patients provided written informed consent.

Gem Hunters is a three-armed probabilistic reward task in which stimuli (gems) vary along two dimensions: shape and color. Each game (6 games; 18 trials/game) contains one relevant dimension (e.g., shape) and one target feature within that dimension (e.g., circle) associated with a higher reward probability (80/20). At the start of each game, patients are explicitly informed of the relevant dimension, thereby providing an efficient state representation *a priori*. Adaptive behavior thus requires selectively attending to the relevant dimension and learning, through trial and error, which of its features is most likely to yield reward. Patients were instructed to respond as quickly as possible and were aware that each trial would time out after 20 s. Reward outcome was presented on screen for 1.5 s immediately following choice. If patients were rewarded, “+1” was depicted onscreen. If patients were not rewarded, “+0” was depicted onscreen. Reward presentation was followed by an inter-trial interval (ITI) in which the screen was blank. To avoid bleeding effects in the neural signal, the ITI period was sampled randomly from a uniform distribution and lasted between 0.5 and 1.5 s before the next trial commenced.

Target features change across games and are signaled to patients. Feature selection is randomized such that each of the six possible target features appears once; patients are not informed of this structure. Before testing, patients viewed standardized instructional videos (available in the software repository) and completed at least one practice game. During the experiment, patients earned a performance-based bonus of $0.03 for each correct choice (i.e., selecting the stimulus containing the target feature), in addition to baseline compensation.

### LFP data recording and preprocessing

Local field potential (LFP) data were recorded from twenty-two patients with drug-resistant epilepsy (DRE) undergoing intracranial monitoring at Mount Sinai West Hospital. Electrode implantation sites were determined solely on clinical grounds by the treating neurosurgical team. All research procedures were approved by the Institutional Review Board of the Icahn School of Medicine at Mount Sinai, and all patients provided written informed consent prior to participation.

Intracranial signals were acquired using the Natus NeuroWorks clinical monitoring system at a sampling rate of 1,500 Hz. Depth electrodes (Ad-Tech Medical) consisted of multi-contact platinum–iridium contacts (1.3 mm diameter, 5 mm spacing) implanted stereotactically along clinically determined trajectories.

For analysis, data were down sampled to 500 Hz and notch-filtered to remove line noise at 60 Hz and its harmonics. Signals were re-referenced in a bipolar configuration to minimize volume conduction. Periods containing interictal epileptiform activity or excessive noise were automatically identified and excluded using a previously validated detection algorithm^54^. On average, a small proportion of data was removed across patients (mean ± SD: OFC = 0.10 ± 0.01; LPFC = 0.13 ± 0.12). Time frequency decomposition was performed using complex Morlet wavelets with logarithmically spaced center frequencies, allowing for constant relative bandwidth across the analyzed frequency range. Spectral power was normalized by z-scoring each epoch (−1.5 to +1.5 s around choice/reward) for each bipolar derivative relative to a 0.5-s baseline epoch corresponding to the inter-trial interval (ITI), during which a blank screen was presented. The baseline ITI immediately preceded the experimental epoch, a conservative approach that minimizes inflation of z-score values. Normalization was performed after automated detection and exclusion of interictal epileptiform discharges and other noise artifacts, with omissions applied to both baseline and experimental epochs prior to frequency decomposition to prevent bias in normalized power estimates. To correct for temporal offsets between the behavioral and neural recording systems, we implemented an automated synchronization procedure that aligned behavioral event timestamps from the task computer with photodiode pulses recorded on a dedicated Natus channel marking key trial events. All preprocessing and analyses were conducted in Python using MNE-Python^36^ and custom code available in the project’s software repository.

### Single-unit data recording and preprocessing

Single-unit activity was recorded from five patients who, in addition to clinical macroelectrodes, were implanted with Behnke–Fried depth electrodes (Ad-Tech Medical) containing microwire bundles. Patients provided informed consent for the implantation of research-only microwires. Implantation procedures and data acquisition followed previously described protocols and were approved by the Icahn School of Medicine at Mount Sinai’s Institutional Review Board where all data collection occurred.

Each Behnke–Fried electrode contained eight 40-µm platinum–iridium microwires extending from the electrode tip and targeted regions selected for research purposes. Neural signals were recorded at 30 kHz using Neuralynx recording systems. Spike detection and sorting were performed using Combinato^55^. Units with mean firing rates below 0.10 Hz or above 10 Hz were excluded. Putative single-units were manually confirmed based on established criteria (Supplementary Fig. 1B). For each retained unit, we computed a time-resolved firing rate during the experimental epoch of interest (−1.5 to 1.5 s around choice/reward) using a sliding window (250 ms) across the spike train. Firing rates were normalized by subtracting the mean firing rate during the baseline inter-trial interval (0.5 s), isolating task-evoked changes in activity.

### Anatomical localization of macro- and microwire electrodes

The anatomical location of each implanted electrode contact was determined using a combination of pre-implantation T1-weighted magnetic resonance imaging (MRI) and post-implantation computed tomography (CT) scans. Co-registration and localization were performed using LeGUI^56^, an open-source MATLAB-based graphical user interface for electrode localization in intracranial electrophysiology.

Briefly, pre-implantation MRIs were first aligned to the post-implantation CT using a rigid-body transformation implemented in SPM12. The CT was then co-registered to the patient’s MRI in native space to visualize electrode contacts relative to individual cortical anatomy. LeGUI^56^ automatically detected electrode contacts based on CT intensity thresholds, and the resulting coordinates were manually inspected and corrected when necessary to ensure alignment along the expected electrode trajectory. Electrode coordinates were then transformed into MNI (Montreal Neurological Institute) standard space using nonlinear normalization of the T1-weighted MRI to MNI152 space. Anatomical labels were assigned by overlaying the transformed electrode coordinates onto the Yale Brain Atlas^57^, a parcellation specifically developed from iEEG studies and optimized for human electrophysiological research^57^.

For microwire bundles, the location of each Behnke–Fried depth electrode was determined by the position of the macroelectrode shaft tip from which the microwires extended. Macroelectrode and microwire locations were visualized in both subject-native and MNI space for verification. Localization results were exported as volumetric and tabular outputs from LeGUI^56^ for subsequent ROI–based analyses.

### Statistical analysis

No statistical methods were used to predetermine sample sizes, but our sample sizes are comparable to those reported in previous publications. For the linear encoding models and cluster detection analyses described in this study, we used a non-parametric permutation method to generate many permutations in which observations are permuted with respect to the null hypothesis. This enabled us to determine critical statistics and p-values (permutation adjusted) against empirically derived null distributions. Where appropriate, parametric statistics were leveraged for hypothesis testing.

### Behavioral modeling of selective attention during multidimensional RL

Following task completion, patients’ observable data included their trial-by-trial choices, received rewards, and the stimuli encountered. However, internal cognitive variables, such as top-down attentional focus, are not directly observable to the experimenter. To infer whether patients deployed top-down, selective attention during multidimensional RL and to formalize these latent processes, we developed two attention-aware, feature-based RL models^7,8^. Both models implemented a Rescorla–Wagner learning rule^58^, wherein a single RPE is computed and used to update feature values on each trial. Importantly, the models differ only in how they allocate attention across stimulus dimensions. Each assumes that patients learn independent feature values (e.g., the value of “orange” and the value of “square”) that are subsequently combined to guide choice, rather than learning a single, conjunctive value for each multidimensional stimulus. In the Uniform Attention RL (UA-RL) model, attention is distributed equally across the two stimulus dimensions (shape and color) during choice and value updating. The Selective Attention RL (SA-RL) model implements selective attention to the instructed relevant dimension, biasing both choice and value updating in favor of the instructed relevant dimension.

In both models, we assume patients choose between available stimuli based on their EV, which is a linear combination of associated features scaled by a selective attention weight:

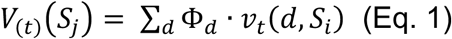

*V*_(*t*)_(*S*_j_) is the value of stimulus *j* on trial *t*, Φ_*d*_ is the selective attention weight on dimension *d*, and *v*_*t*_(*d*, *S*_*i*_) denotes the value of the feature in dimension *d* of stimulus *S*_*i*_. Following feedback, an RPE is calculated:

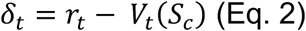

Where *V*_*t*_(*S*_*c*_) is the chosen stimulus’ EV. The RPE updates the chosen stimuli’s associated feature values:

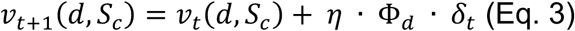

The update is scaled by learning rate *η*. Choice probability was computed using a softmax action selection rule. The SA-RL model’s Φ is a free parameter implementing selective attention to favor the relevant dimension (Eq.1/3). The UA model’s Φ is fixed at 0.50 for both dimensions, thus implementing uniform attention across dimensions during decision-making and value updating.

Critically, the SA-RL model extends previous work^7,8^ by fitting selective attention as a subject-level free parameter, rather than as a trial-by-trial dynamic variable. This approach yields subject-specific estimates of attentional focus that reflect stable individual differences in the maintenance of an efficient state representation. By treating attention as a static, trait-like parameter, we capture variability in patients’ strategies during task completion. These fitted attention parameters can then be related to individual differences in neural encoding and connectivity, providing a direct link between behavioral strategy and underlying neural mechanisms.

### Behavioral model validation, fitting, and evaluation

To assess how well each model captured patients’ choice behavior, we first conducted parameter and model recovery analyses (Supplementary Fig. 2). This procedure involved simulating synthetic datasets across a broad range of parameter values for each model, refitting the models to these data, and verifying that the recovered parameters and model identities matched the generative ones. Model fitting during recovery used the same optimization and cross-validation procedures as those applied to the empirical data. Both models exhibited high recoverability, indicating that their parameters were identifiable and that the models could be reliably distinguished. Specifically, the SA-RL model was correctly recovered in 96% of simulations, whereas the UA-RL model was recovered in 66% of simulations, with both rates exceeding chance (Supplementary Fig. 2C). Parameter recovery analyses further confirmed robust identifiability. For the SA-RL model, recovered and true parameters were strongly correlated for the learning rate (Pearson r = 0.83, p < 0.001) and attention weight (r = 0.82, p < 0.001), with a moderate correlation for the inverse temperature (r = 0.28, p < 0.01; Supplementary Fig. 2A). For the UA-RL model, parameter recovery was strong for the learning rate (r = 0.85, p < 0.001) and moderate for the inverse temperature (r = 0.41, p < 0.001; Supplementary Fig. 2B).

We next fit each model to patients’ trial-by-trial choice data using leave-one-game-out cross-validation with maximum likelihood estimation. For each patient, models were trained on five games and tested on the held-out sixth game; this procedure was repeated such that each game served as the test set once. To mitigate local minima, model fitting was repeated for five iterations using random starting values for each free parameter. For each iteration, predictive log likelihoods were averaged across test games, and the parameter set yielding the highest average test likelihood was retained as the best-fitting solution for those patients.

Model optimization was performed in R using the optim function with the L-BFGS-B algorithm. Model performance was assessed by comparing the average predictive log likelihood of observed choices under each fitted model. The model with the higher predictive likelihood was taken to provide the better account of behavior. All model code and fitting procedures are publicly available in the project’s software repository.

### Cluster-based permutation testing of task-evoked time-frequency activity

To assess whether task engagement elicited significant modulation of oscillatory activity relative to baseline, we performed non-parametric cluster-based permutation tests^36^ on time frequency representations of LFP power across frequencies. Time-frequency representations of normalized power were pooled across electrodes within each region, and a one-sample cluster-based permutation test was applied to identify time frequency clusters where power significantly deviated from baseline at the group level. Clusters were formed by identifying contiguous time–frequency points exceeding a predefined threshold and summing the associated test statistics within each cluster. Statistical significance was determined by comparing observed cluster statistics to a null distribution generated by randomly sign-flipping data across permutations, thereby controlling for multiple comparisons across time and frequency. This approach enabled identification of task-evoked oscillatory activity without assumptions about the temporal or spectral extent of effects.

### Electrode-level trial-resolved regression analysis of EV encoding

To examine how value-related variables were encoded in LFP activity, we used a multivariate linear encoding framework applied separately to HFA (60–200 Hz) and theta-band power (3–8 Hz). For each electrode, band-limited power time series were computed and averaged across trials within a −1.5 to +1.5 s window surrounding choice and reward outcome. The resulting trial × power matrices served as input to electrode-level encoding models relating behaviorally derived regressors to neural activity. Separate multivariate linear models were fit for each electrode. Unless otherwise noted, all modeling procedures were identical for HFA and theta-band analyses.

We employed linear encoding models because they are computationally efficient, yield interpretable parameters, and are well suited for hypothesis-driven tests of whether cognitive variables such as expected value (EV) predict neural activity. For each electrode, we generated a within-electrode null distribution by shuffling the regressor of interest across trials (1,000 iterations) and refitting the model; observed regression coefficients were then z-scored relative to this null distribution.

Within this encoding framework, we first asked which value-related variable best explained local neural activity in each region. We compared the average log-likelihoods of multivariate models predicting HFA that each included a single value-based regressor, EV, reward, or reward prediction error (RPE), alongside value-agnostic task features (chosen shape, chosen color, and the relevant dimension; Supplementary Table 3). The model including EV achieved higher average log-likelihoods across electrodes than models including reward or RPE.

This empirical result was complemented by a theoretical motivation for focusing on EV. Unlike reward or RPE, EV reflects a subjective value estimate that depends on the internal state representation maintained by the patients. Because the selective attention parameter compresses feature values in a task-relevant manner, EV directly indexes the attention-weighted state representation that guides behavior. Moreover, EV is uniquely available both before and after choice, enabling a unified analysis of anticipatory and outcome-related neural dynamics. For these reasons, EV provided both the strongest explanatory power and the most theoretically relevant variable for probing state representations in prefrontal cortex.

The final equation for the electrode-level regression was:

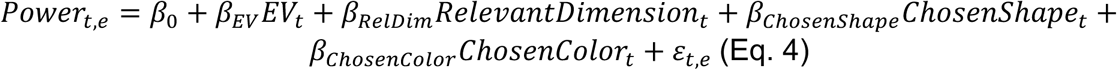

Where *Power*_*t*,*e*_ denotes either HFA or theta power on trial *t* for electrode *e*. *β*_0_ is the intercept, the predictors encode trial-wise EV, the current relevant dimension (shape or color), and the categorical feature-level identity of the chosen shape and color, and *ɛ*_*t*,*e*_ is the residual error term. Electrode-wise EV effects (z-scored *β*_*EV*_ estimates) were derived using a permutation-based procedure in which the EV regressor was randomly shuffled across trials, the GLM refit, and a null distribution of EV effect sizes computed for each electrode; the observed *β*_*EV*_ was then standardized relative to this electrode-specific null. To account for unequal electrode sampling across patients, we used an intercept-only mixed-effects model to aggregate electrode-wise z-scored effects and assess whether population-level encoding for each regressor differed significantly from zero. This two-level approach preserves electrode-specific signal structure while appropriately modeling subject-level variability. Electrode-wise EV effects (z-scored *β*_*EV*_ estimates) were then entered into a linear mixed-effects model to evaluate whether EV encoding was consistently positive across patients while accounting for differences in electrode count. The model was:

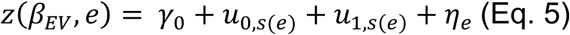

This corresponds to the formula:

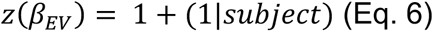

Where *z*(*β*_*EV*_, *e*) is the standardized EV effect size for electrode *e*_*t*_, *γ*_0_ is the population level intercept, *u*_0,*s*(*e*)_ is the subject-specific random intercept, *u*_1,*s*(*e*)_ is the subject-specific random slope (accounting for variation in the number of electrodes across patients), and *η*_*e*_ is the residual error.

To assess the relationship between selective attention (Φ) and value encoding, we regressed the EV effect size for each electrode against the corresponding subject-specific selective attention weight (Φ), separately for HFA and theta-band activity.

To evaluate encoding of control regressors in each model (Eq. 4), we repeated the permutation procedure while shuffling each regressor independently, thereby disrupting its specific association with the neural data while preserving all other relationships among variables. Group-level effects were assessed using an intercept-only model (see Results). Electrode-wise regression coefficients for all regressors are reported in Supplementary Fig. 4.

### Time-resolved regression of EV encoding across regions

To characterize the temporal dynamics of EV encoding in LPFC and OFC, we extended the electrode-level multivariate regression analysis into the temporal domain by performing a time-resolved regression at each time point across the experimental epoch (−1.5 to +1.5 s relative to choice and outcome). At each time point, regression coefficients were estimated independently, capturing the instantaneous relationship between neural activity and task variables. The same set of predictors used in the time-averaged models was included at each time point. HFA and theta-band power were analyzed separately using identical procedures. Specifically, neural power was modeled as a function of the EV of the chosen option, while controlling for the relevant dimension and chosen feature. The resulting time-resolved EV-related β coefficients quantified the moment-by-moment strength of value encoding for each electrode across the trial.

To identify time windows within each region showing significant EV-related effects, we performed a one-sample permutation-based cluster test^36^ on electrode-level *β* coefficients against zero. Temporally adjacent significant samples were clustered, and cluster statistics were evaluated against a permutation-derived null distribution to control for multiple comparisons across time.

To further characterize the temporal structure of value encoding at the electrode level, we next quantified the latency of peak EV encoding for each electrode, defined as the time point within the decision epoch at which the absolute EV-related regression coefficient reached its maximum. Distributions of peak encoding times were then computed separately for each frequency band and region. To test for differences in the distribution of peak encoding times between selective attention groups (median split of Φ), we used a nonparametric two-sample Kolmogorov–Smirnov test. All temporal analyses were performed identically for HFA and theta-band activity.

### Detection of theta-band oscillations

To assess the presence of theta-band oscillations, we parameterized electrode-wise power spectra using the FOOOF algorithm, which separates periodic oscillatory components from the aperiodic (1/f-like) background of neural power spectra^59^.

Power spectral densities were computed for each electrode using a multitaper approach and averaged across trials to obtain a single spectrum per electrode. Spectra were fit over the 1–30 Hz frequency range using FOOOFGroup with parameters matched to those used by Donoghue and colleagues^59^ (peak width limits of 1–8 Hz, minimum peak height of 0.1 (relative to the aperiodic fit), peak threshold of 1, and a maximum of three peaks per spectrum).

Theta-band oscillations were defined as periodic peaks with center frequencies between 3 and 8 Hz. For electrodes exhibiting multiple theta-band peaks, we selected the peak with the highest relative power. For each electrode, we extracted the theta peak’s center frequency, relative power, and the number of detected theta peaks.

Electrodes were assigned to anatomical regions of interest (OFC, LPFC) based on subject-specific electrode localization. For each region, theta peak power values were aggregated across electrodes and patients.

To determine whether theta-band oscillations were reliably present at the group level, we used the same modeling strategy applied to assess group-level encoding of cognitive regressors. Specifically, we fit an intercept-only linear mixed-effects model to theta peak power values within each region, with subject included as a random effect. Significance of the intercept was assessed to test whether mean theta power differed from zero across patients (OFC: *β* = 0.24, z= 10.38, *p* < 0.001; LPFC: *β* = 0.39, z= 7.18, *p* < 0.001). This analysis provided a conservative criterion for confirming that regions of interest exhibited reliable theta-band oscillations across patients (Supplementary Fig. 5).

### Computation of theta-band directed functional connectivity between OFC and LPFC

To quantify directed functional connectivity, we used the Phase Slope Index (PSI), a frequency-resolved metric that infers the direction of information flow by detecting consistent phase leads across neighboring frequencies. PSI was computed using MNE-Python’s implementation^36^. Because PSI is estimated across trials, we examined value-related differences in connectivity by splitting trials into high- and low-expected value (EV) conditions via a mean split of trial-wise EVs. For each unilateral LPFC–OFC electrode pair, we computed time-resolved PSI in the theta band (3–8 Hz) separately for the two conditions and then averaged across pairs.

Statistical significance of EV-related differences in connectivity dynamics was assessed using a time-resolved, cluster-based two-sample t-test^36^. Guided by prior evidence implicating the LPFC in top-down attentional control and our own findings that LPFC local value encoding is attention-dependent, we adopted the framework that information flow originates in LPFC and projects to OFC. Accordingly, positive PSI values indicate the LPFC leads the OFC, whereas negative values indicate the OFC leading the LPFC. This approach enables us to capture the temporal evolution of directed, frequency-specific connectivity between these regions.

### Relating LPFC HFA value signals to LPFC–OFC EV-based connectivity

To relate LPFC HFA value signals to LPFC–OFC EV-dependent connectivity, we computed time-resolved differences in phase slope index (PSI)^36^ between high- and low-EV trials within a 500 ms pre-choice window. This window encompassed the period of significant EV-based modulation of LPFC–OFC theta-band connectivity, as identified using a two-sample cluster-based permutation test^36^ (Fig. 6). For each time point, we defined ΔPSI as the difference between PSI estimates on high- and low-EV trials (ΔPSI = PSIhigh EV − PSIlow EV). Under this convention, positive ΔPSI values indicate stronger LPFC-to-OFC directed connectivity on high-EV relative to low-EV trials, whereas negative values indicate stronger LPFC-to-OFC connectivity on low-EV trials. This approach yielded a single time-resolved metric capturing the magnitude and direction of EV-dependent modulation of LPFC–OFC connectivity, which was subsequently related to LPFC HFA value encoding.

For local LPFC HFA activity, we extracted time-resolved EV-related effect sizes, defined as the regression coefficients (β) from the time-resolved encoding analysis described previously (Fig. 4C). Both the HFA EV-related coefficients and the LPFC–OFC connectivity measures were restricted to the same 500 ms pre-choice window to enable direct temporal correspondence between local value encoding and inter-regional connectivity.

We fit a linear mixed effects model predicting ΔPSI (high–low EV) from LPFC EV encoding (HFA∼EV β), its interaction with selective attention weight (Φ), and random effects accounting for between-electrode variability. The model was:

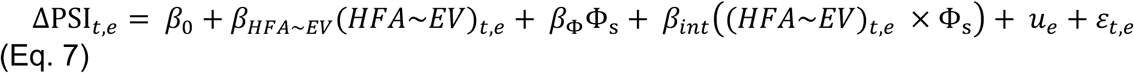

Here, ΔPSI_*t*,*e*_ denotes the difference in LPFC–OFC theta-band directed connectivity between high- and low-EV trials at time point *t* for electrode *e*. *β*_0_ is the fixed intercept, and fixed-effect predictors include the strength of local LPFC value encoding, quantified as the time-resolved HFA–EV regression coefficient (*HFA*∼*EV*)_*t*,*e*_, the subject-level selective attention parameter Φ_s_, and their interaction. A random intercept *u*_*e*_ accounts for between-electrode variability, and *ɛ*_*t*,*e*_ denotes the residual error term. This model tests whether EV-dependent modulation of LPFC–OFC connectivity is predicted by local LPFC value signals and whether this relationship is systematically shaped by individual differences in selective attention.

### Single-unit trial-resolved regression analysis

We applied the same EV encoding model used for the LFP analyses (Eq. 4; Supplementary Table 3), using EV, the relevant feature dimension, and the chosen stimulus features to predict trial-resolved mean firing rate. For each putative single-unit, we fit a non-parametric multivariate linear encoding model to quantify how these task variables predicted spiking activity on a trial-by-trial basis. To assess significance, we constructed a permutation-derived null distribution for each regressor by shuffling that regressor across trials and recomputing the encoding model. Observed regressions coefficients were standardized relative to this unit-specific null distribution. This procedure allowed us to identify units tuned to EV, to the relevant dimension, or to both, and to quantify the proportion of neurons selective for each state-relevant variable.

## Code Availability

Analysis code will be made available in an open-source repository upon manuscript acceptance.

