## Supplemental Information for "Coordinated human prefrontal dynamics sustain task-state representations during learning"

### Supplementary Information

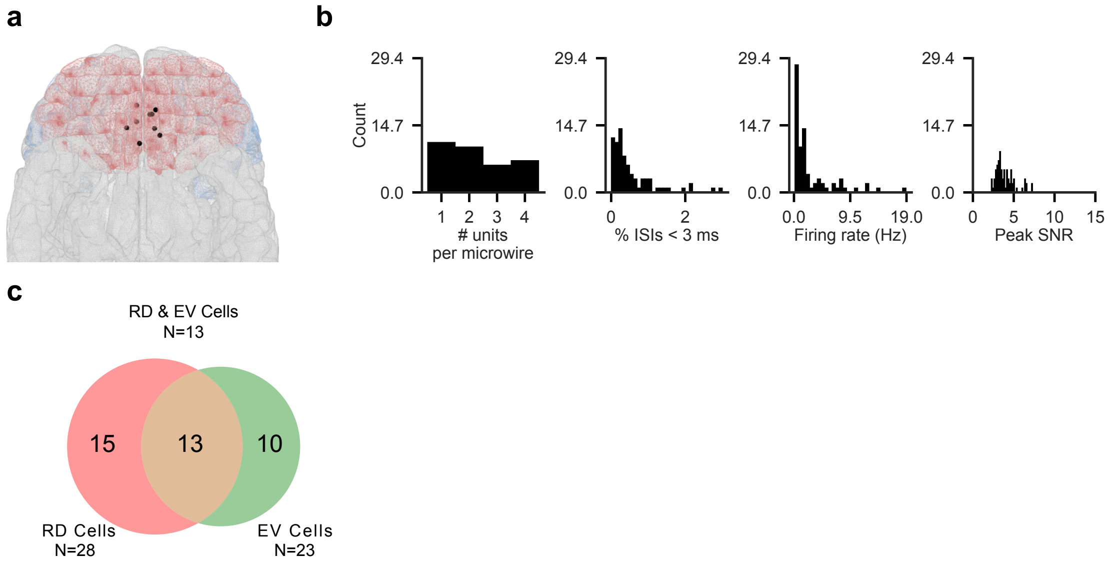


**Supplementary Figure 1. Single unit recordings. A.** Locations of microelectrode bundle placements in OFC (N = 8 bundles) across five patients who consented to microelectrode recordings. **B.** Following preprocessing and manual spike sorting, 72 putative single units met standard quality criteria and were retained for analysis; distributions of quality metrics for retained units are shown. **C.** Venn diagram illustrating the proportion of units selectively tuned to EV, the task-relevant dimension, or both.

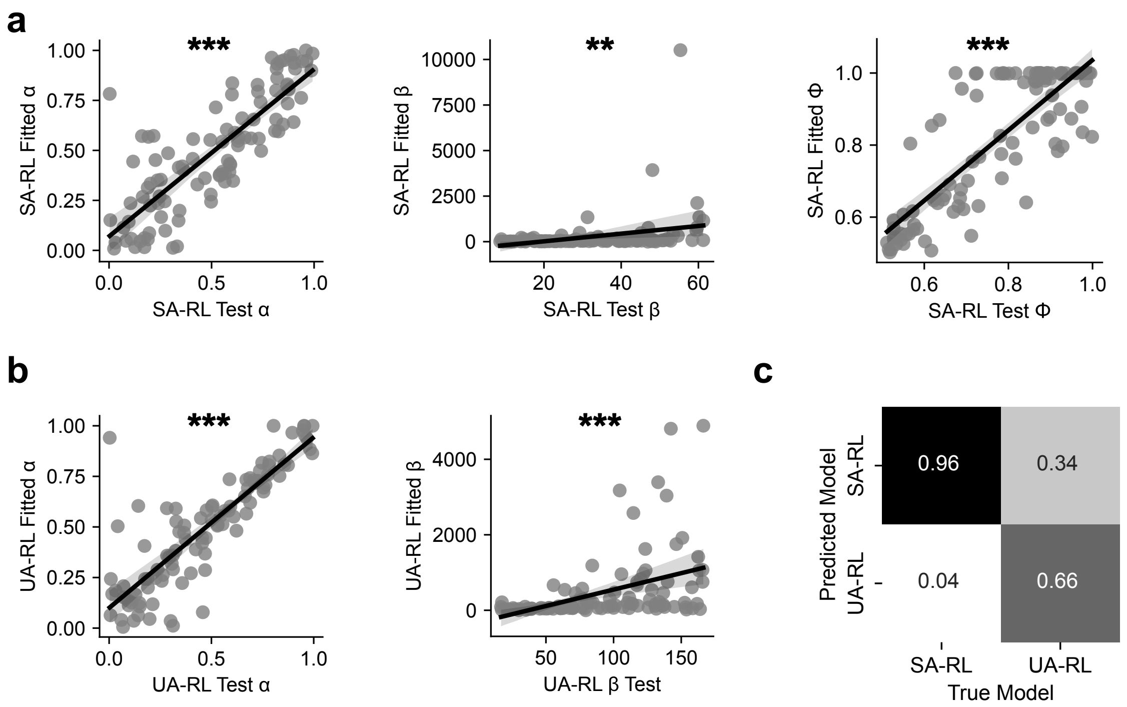


**Supplementary Figure 2. Parameter and model recovery for behavioral models. A.** Parameter recovery for the SA-RL model's free parameters (left to right: learning rate α, inverse temperature β, and selective attention weight Φ). We simulated 100 datasets using randomly sampled parameter values within the fitting bounds and refit the model to each dataset. Recovered parameters closely matched the generative values for all parameters, indicating reliable parameter recovery Model recovery and parameter identifiability were high (SA-RL: learning rate, $r = 0.83$, $p < 0.001$; attention weight, $r = 0.82$, $p < 0.001$; inverse temperature, $r = 0.28$, $p < 0.01$; UA-RL: learning rate, $r = 0.85$, $p < 0.001$; inverse temperature, $r = 0.41$, $p < 0.001$). B. Parameter recovery for the UA-RL model's free parameters (left to right: learning rate α and inverse temperature β), assessed using the same simulation and refitting procedure. All parameters showed strong recovery. **C.** Model recovery analysis. For 100 simulated datasets generated from each model, we fit both models and identified the model with the higher average predictive log likelihood as the best-fitting model. Results are summarized in a confusion matrix, showing that most datasets for each model were correctly assigned to their generative model, indicating reliable model identifiability (model recovery: SA-RL, 96%; UA-RL, 66%, both > chance).

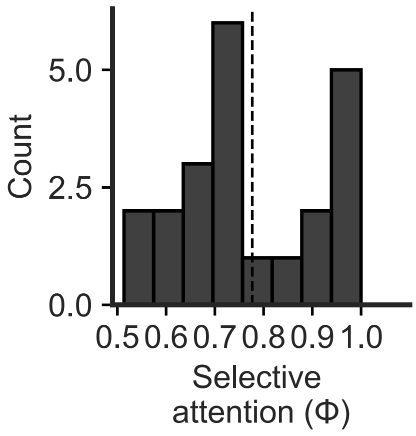


**Supplementary Figure 3. Distribution of selective attention estimates from the SA-RL model.** Shown is the distribution of subject-level estimates of the selective attention parameter (Φ) from the SA-RL model across participants (N = 21; mean ± SD = 0.78 ± 0.15). Parameter estimates were obtained using leave-one-game-out cross-validation with maximum-likelihood estimation of each participant's sequence of choices under the SA-RL model (see Methods).

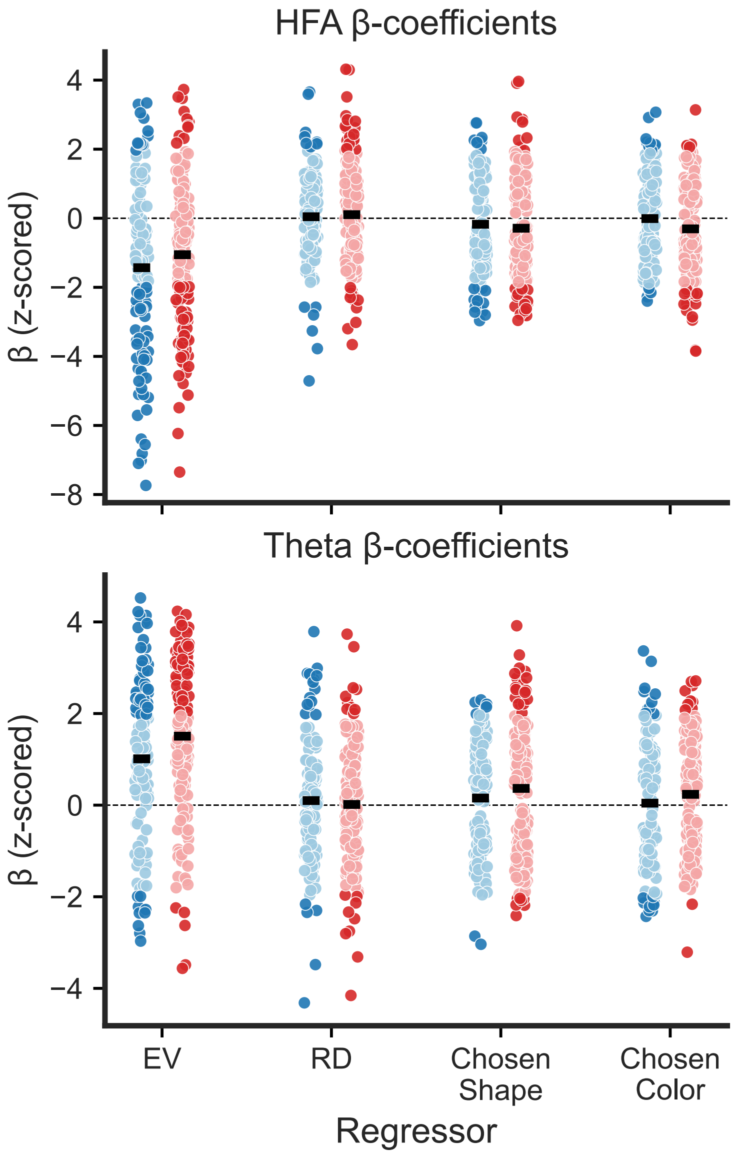


**Supplementary Figure 4. Electrode-level β-coefficients for trial-resolved regression**. Scatter plots show permutation-based, z-scored regression coefficients (β) for individual electrodes in LPFC (blue) and OFC (red) for EV, relevant dimension (RD), chosen shape, and chosen color. Each point corresponds to a single electrode; darker colors indicate electrodes with significant effects (|z| > 1.96), and lighter colors indicate non-significant electrodes. Horizontal black bars denote the mean z-score within each region and regressor. The top panel shows HFA, and the bottom panel shows theta-band activity. Dashed horizontal lines indicate zero. Proportion of significant electrodes in HFA (top; LPFC: EV = 0.49, RD = 0.11, Chosen Shape = 0.16, Chosen Color = 0.13; OFC: EV = 0.40, RD = 0.19, Chosen Shape = 0.19, Chosen Color = 0.17). Proportion of significant electrodes in theta (bottom; LPFC: EV = 0.43, RD = 0.16, Chosen Shape = 0.06, Chosen Color = 0.19; OFC: EV = 0.44, RD = 0.12, Chosen Shape = 0.18, Chosen Color = 0.07).

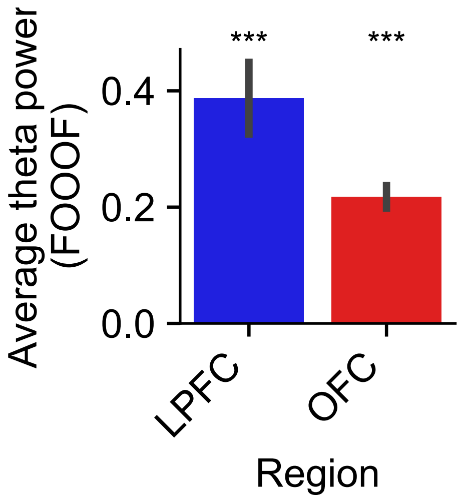


**Supplementary Figure 5. Detection of theta oscillations using FOOOF.** Theta-band power was extracted on a per-electrode basis from the periodic component of FOOOF fits to LFP power spectra in LPFC, OFC. To account for unequal electrode sampling across patients, an intercept-only mixed-effects model with subject as a random effect was fit for each region, testing whether mean theta power was significantly greater than zero at the group level. Both regions exhibited theta power significantly above zero (OFC: $\beta$ = 0.24, z= 10.38, $p$ < 0.001; LPFC: $\beta$ = 0.39, z= 7.18, $p$ < 0.001), confirming the presence of robust theta oscillations across the sample and motivating subsequent theta phase–based connectivity analyses.

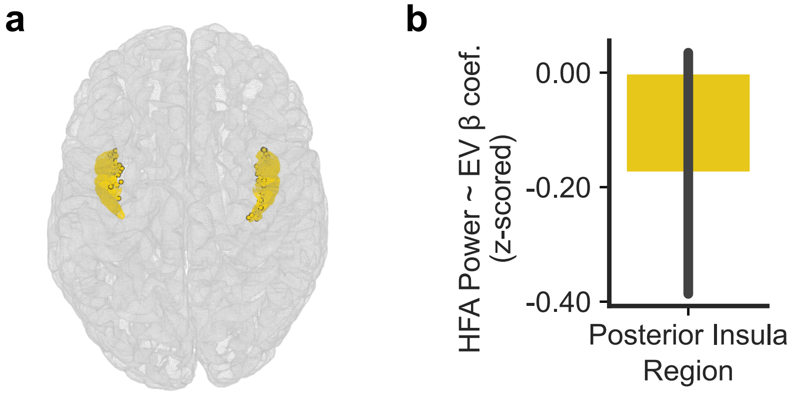


**Supplementary Figure 6. Posterior insula as a control region for EV encoding. A.** Locations of macroelectrodes in posterior insula, which was included as a control region ($N$ = 61 electrodes). **B.** Trial-resolved encoding analyses revealed no consistent encoding of EV in posterior insula across patients (β = -0.35, $z$ = -1.05, $p$ > 0.05). In contrast to LPFC and OFC, EV-related HFA effects in posterior insula did not show a reliable group-level pattern.

**Table 1. Participant Demographics**

| | |
|---|---|
| Age (M ± SD) | 39.2 ± 13.8 |
| Sex | 15 F / 7 M; no other identities reported |
| Race | 41% White |
| Ethnicity | 55% non-Hispanic |
| Education Level | 41% bachelor's degree (or higher) |

**Supplementary Table 1. Demographic information.** Summary of demographic characteristics for the study's patients.

**Table 2. Counts of LPFC and OFC Bipolar LFP Derivatives and OFC Single Units per Subject**

| Subject | LPFC LFP | OFC LFP | OFC Single Units |
|---|---|---|---|
| 1 | 9 | 5 | N/A |
| 2 | 6 | 4 | N/A |
| 3 | 1 | 5 | N/A |
| 4 | 3 | 7 | N/A |
| 5 | 5 | 6 | N/A |
| 6 | 4 | 4 | N/A |
| 7 | 1 | 4 | N/A |
| 8 | 2 | 6 | N/A |
| 9 | 3 | 9 | 9 |
| 10 | 1 | 14 | 22 |
| 11 | 1 | 15 | N/A |
| 12 | 6 | 12 | 16 |
| 13 | 3 | 4 | 11 |
| 14 | 3 | 8 | N/A |
| 15 | 12 | 10 | N/A |
| 16 | 15 | 17 | N/A |
| 17 | 7 | 1 | N/A |
| 18 | 2 | 14 | N/A |
| 19 | 5 | 2 | N/A |
| 20 | 7 | 4 | N/A |
| 21 | 15 | 8 | N/A |
| 22 | 11 | 4 | 14 |
| Total | 122 | 163 | 72 |

**Supplementary Table 2. Electrode coverage.** Number of bipolar LFP derivatives recorded in LPFC and OFC, and number of well-isolated OFC single units retained for analysis for each subject. Totals across patients are reported in the final row. All patients contributed at least one macroelectrode in both LPFC and OFC. Five patients also consented to microelectrode implantation; NA denotes patients without microelectrode recordings.

**Table 3. Regional Log-Likelihoods for Value-Based HFA Encoding Models**

| Model | Region | Log Likelihood |
|---|---|---|
| **EV model** (EV + Relevant Dimension + Chosen Shape + Chosen Color) | OFC | 104.2 |
| | LPFC | 90.5 |
| **RPE model** (RPE + Relevant Dimension + Chosen Shape + Chosen Color) | OFC | 102.2 |
| | LPFC | 86.7 |
| **Reward model** (Reward + Relevant Dimension + Chosen Shape + Chosen Color) | OFC | 102.5 |
| | LPFC | 87.6 |

**Supplementary Table 3. Model comparison of value-related predictors for HFA encoding.** Average log likelihoods across electrodes in OFC and LPFC for multivariate encoding models predicting HFA. For each electrode, separate models were fit that included one value-related regressor, EV, RPE, or reward, along with state-related covariates (relevant dimension, chosen shape, and chosen color). Values shown reflect the mean log likelihood across electrodes within each region.
